# Myeloid-biased HSC require Semaphorin 4A from the bone marrow niche for self-renewal under stress and life-long persistence

**DOI:** 10.1101/2022.01.12.476097

**Authors:** Dorsa Toghani, Sharon Zeng, Elmir Mahammadov, Edie I. Crosse, Negar Seyedhassantehrani, Christian Burns, David Gravano, Stefan Radtke, Hans-Peter Kiem, Sonia Rodriguez, Nadia Carlesso, Amogh Pradeep, Nicola K. Wilson, Sarah J. Kinston, Berthold Göttgens, Claus Nerlov, Eric Pietras, Marion Mesnieres, Christa Maes, Atsushi Kumanogoh, Thomas Worzfeld, Peter Kharchenko, David T. Scadden, Antonio Scialdone, Joel A Spencer, Lev Silberstein

## Abstract

Tissue stem cells are hierarchically organized. Those that are most primitive serve as key drivers of regenerative response but the signals that selectively preserve their functional integrity are largely unknown. Here, we identify a secreted factor, Semaphorin 4A (Sema4A), as a specific regulator of myeloid-biased hematopoietic stem cells (myHSC), which are positioned at the top of the HSC hierarchy. Lack of Sema4A leads to exaggerated myHSC (but not downstream “balanced” HSC) proliferation after acute inflammatory stress, indicating that Sema4A enforces myHSC quiescence. Strikingly, aged Sema4A knock-out myHSC expand but almost completely lose reconstitution capacity. The effect of Sema4A is non cell-autonomous, since upon transplantation into Sema4A-deficient environment, wild-type myHSC excessively proliferate but fail to engraft long-term. Sema4A constrains inflammatory signaling in myHSC and acts via a surface receptor Plexin-D1. Our data support a model whereby the most primitive tissue stem cells critically rely on a dedicated signal from the niche for self-renewal and life-long persistence.

## INTRODUCTION

In multiple tissues, stem cells are hierarchically organized and contain distinct, functionally specialized subsets. For example, in the brain (Sachewsky et al., 2019), skeletal muscle (Scaramozza et al., 2019), cornea (Altshuler et al., 2021; Farrelly et al., 2021) and skin (Hsu et al., 2011; Rompolas et al., 2013), the most primitive stem cells (which are also largely quiescent) become activated by injury or stress, whereas their “downstream”, more differentiated counterparts are more proliferative and mainly engaged in on-going tissue repair. Although this two-tiered organization of the stem cell compartment is essential for life-long tissue maintenance, the mechanisms that ensure functional preservation and persistence of individual stem cell subsets within a tissue stem cell hierarchy remain largely unknown.

In the bone marrow, the stem cell hierarchy is exemplified by the myeloid-biased and “balanced” (HSC) subsets, in which the former is considered the most primitive (Challen et al., 2010; Morita et al., 2010; Sanjuan-Pla et al., 2013). Compared to balanced HSC (balHSC), myeloid-biased HSC (myHSC) are inherently skewed towards myeloid differentiation, endowed with a higher self-renewal potential and possess a superior ability to enter cell cycle in response to inflammatory stress (Mann et al., 2018; Matatall et al., 2014; Mitroulis et al., 2018). Although these properties are beneficial for powerful and timely host defense response, they likely account for myHSC expansion during inflammation and aging, which is associated with profound and irreversible functional loss (Beerman et al., 2010; Esplin et al., 2011; Grover et al., 2016; Pang et al., 2011). Thus, despite being positioned at the top of the HSC hierarchy, myHSC appear most vulnerable to stress-induced damage.

In the current study, we identify a membrane-bound and secreted protein Semaphorin 4A (Sema4A) as a myHSC-protective factor. We show that under stress conditions, such as aging and transplantation, the absence of Sema4A results in excessive expansion and functional attrition of myHSC. Surprisingly, balHSC are only minimally affected, suggesting that the effect of Sema4A is myHSC-specific. We further demonstrate that Sema4A from the bone marrow niche is essential for myHSC self-renewal and identify Plexin-D1 as a functional receptor. Our results reveal that by selectively preserving a functional myHSC pool, Sema4A plays a key role in long-term maintenance of the HSC hierarchy.

## RESULTS

### Sema4A regulates quiescence of mouse and human hematopoietic stem/progenitor cells

We have previously established proximity-based analysis as a platform for niche factor discovery (Silberstein et al., 2016). We compared single cell RNA-Seq signatures of osteolineage cells (OLC) that were located in close proximity to a single transplanted HSC (proximal OLC) and further away (distal OLC), and functionally validated several secreted factors with a higher expression in proximal OLC (Angiogenin, IL18, Embigin, VEGF-C) as non cell-autonomous regulators of hematopoietic stem cell/progenitor quiescence (Fang et al., 2020; Goncalves et al., 2016; Silberstein *et al.*, 2016). Because Sema4A displayed a similar expression difference by being significantly more abundant in proximal OLC, as shown in Fig. 1A, we hypothesized that it could also act as a niche-derived HSC quiescence regulator.

**Figure 1.**
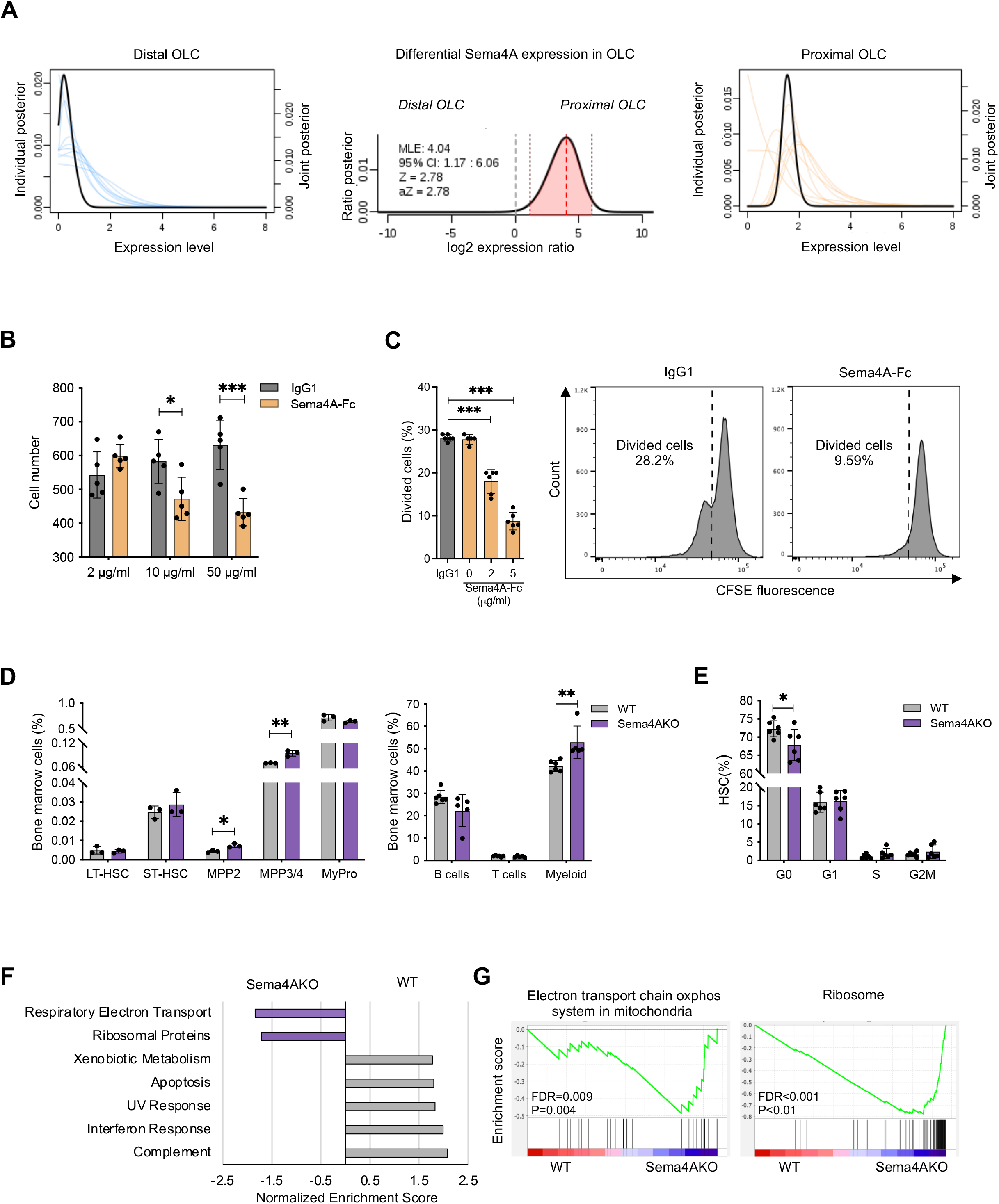
Sema4A regulates quiescence of mouse and human hematopoietic stem/progenitor cells. (A) Expression of Sema4A in single proximal and distal osteolineage cells. The joint posteriors (black lines) describe the overall estimation of likely expression levels within the proximal (top) and distal (bottom) OLCs and are used to estimate the posterior of the expression fold difference (middle plot). The shaded area under the fold-difference posterior shows 95% confidence region. FPM, fragments per million. (B) The number of mouse LKS cells 24 hours after addition of mouse Sema4A-Fc/IgG1 control protein (n=5 technical replicates per condition). (C) CFSE dilution analysis of *ex vivo* proliferation kinetics of human CD34+ cells 24 hours after addition of human Sema4A-Fc/IgG1 protein (n=5 technical replicates per condition, Donor 1). Estimated number of divided cells and representative CSFE fluorescence histograms are shown. (D) Immunophenotypic analysis of the bone marrow from young WT/Sema4AKO mice (n=3-6 per genotype). (E) HSC cell cycle analysis using DAPI/Ki-67 staining in young WT/Sema4AKO mice (n=6 per genotype). (F) Gene set enrichment analysis (GSEA) of single cell RNA-Seq data for the HSC cluster (as defined in Figure S1K) from young WT/Sema4AKO mice. FDR<0.01, top five enriched pathways are shown. (G) GSEA plots for the pathways as shown in (F). Data are presented as mean ± SD *p < 0.05; **p < 0.01; ***p < 0.001 by unpaired T-test.

Semaphorins are a large family of membrane-bound and/or secreted proteins which mediate cell-cell communications in neural development, angiogenesis, immune response and cancer (Fard and Tamagnone, 2021; Kolodkin et al., 1993). In keeping with our hypothesis for a possible non cell-autonomous regulatory role of Sema4A in hematopoiesis, we found that Sema4A transcripts were detectable in niche cell subsets, such as CD31^+^ endothelial cells and niche factor-enriched VCAM1^high^Embigin^+^ OLC fraction, as also supported by the published data (Baccin et al., 2020) (Fig. S1A and S1B). Of note, expression of Sema4A in CD45^−^Ter119^−^ ALCAM^+^ bone-lining cells (Valletta et al., 2020) was significantly increased in aged mice (Fig. S1C). In human bone marrow, Sema4A mRNA was also present in bone-lining and endothelial cells, as demonstrated by single-molecule fluorescent in situ hybridization (Fig. S1D).

In order to gain initial functional insights, we tested the effect of Sema4A on proliferation of mouse and human hematopoietic stem/progenitor cells *in vitro*. Consistent with our hypothesis, addition of recombinant Sema4A-Fc protein resulted in fewer lin^−^c-Kit^+^Sca-1^+^ (LKS) cells after 24 hours of liquid culture and suppression of hematopoietic colony formation in a dose-dependent manner (Fig. 1B and S1E). Similarly, human recombinant Sema4A-Fc inhibited *in vitro* proliferation of human bone marrow CD34^+^ cells from several donors, as measured by the Carboxyfluorescein succinimidyl ester (CFSE) dilution assay (Takizawa et al., 2011) (Fig. 1C and S1F). Collectively, these data indicate that Sema4A non cell-autonomously restricts HSPC proliferation, and that this property is conserved between mice and humans.

Next, we examined the role of Sema4A in steady-state hematopoiesis using a germline knock-out model. Sema4A knock-out (Sema4AKO) mice are viable and have a normal life span (Kumanogoh et al., 2005). However, baseline analysis of young Sema4AKO animals revealed subtle but reproducible anemia and thrombocytosis (Fig. S1G). Moreover, in the bone marrow, while the number and frequency of long-term HSC (LT-HSC, defined as LKS CD48^−^CD34^−^Flk2^−^ CD150^+^) was similar between the genotypes, the differentiation was skewed towards the myeloid lineage, as evidenced by increased frequency of myeloid-committed progenitors MPP2 and mature myeloid cells (Fig. 1D, see Fig. S1H for gating strategy). Importantly, young Sema4AKO HSC displayed more active cycling, as assessed by Ki-67/DAPI staining and EdU incorporation (Fig. 1E, S1I and S1J). Single-cell RNA-Seq of HSPC from young WT and Sema4AKO mice, while showing no difference in cluster distribution (Fig. S1K), revealed positive enrichment for the terms “Kegg ribosome” and “Electron transport chain oxphos system in mitochondria” in Sema4AKO cells within the HSC cluster, consistent with disruption of quiescence (Fig. 1F, 1G, S1L, and Suppl. File). Collectively, these results indicate that loss of Sema4A leads to increased HSC proliferation, metabolic activation and myeloid-biased differentiation.

### Sema4A/PlxnD1 signaling constrains the response of myeloid-biased HSC to proliferative stress

Given recent evidence suggesting that lineage-restricted HSC subsets are differentially regulated by niche-derived signals, we hypothesized that myeloid bias in the Sema4AKO model is due to the lack of quiescence-inducing effect of Sema4A specifically on myHSC. While baseline analysis demonstrated no appreciable difference in cell cycle status between Sema4AKO myHSC (LKS CD48^−^CD34^−^Flk2^−^CD150^high^) and balHSC (LKS CD48^−^CD34^−^Flk2^−^CD150^low^) (Beerman *et al.*, 2010) [data not shown], exposure to acute inflammatory stress revealed important HSC subset-specific differences.

In particular, twenty-four hours after injection with polyinosinic:polycytidilic acid (Poly (I:C)) (Walter et al., 2015)(Fig 2A), we found a significant increase in the percentage of Sema4AKO myHSC in G2M phase of cell cycle (Fig.2B and S2B) while no cell cycle difference was observed in balHSC subset (Fig. 2C and S2C, see Fig. S2A for gating strategy under inflammatory conditions) (Hirche et al., 2017). Of note, myHSC and balHSC cycling was comparable in PBS-injected WT/Sema4AKO animals (Fig S2D and S2E) suggesting that the above changes in Sema4AKO myHSC were due to exaggerated response to inflammatory stress. Indeed, subsequent RNA-Seq analysis of myHSC and balHSC from Poly (I:C)-injected animals revealed enrichment for the terms “IL6-Jak Stat3 signaling” and “Interferon alpha response” which was unique to Sema4AKO myHSC dataset (Fig. 2D, 2E, S2F, Suppl. File). Thus, our results reveal that the absence of Sema4A promotes myHSC cell cycle entry and lead to enhanced myHSC sensitivity to inflammatory signaling.

**Figure 2.**
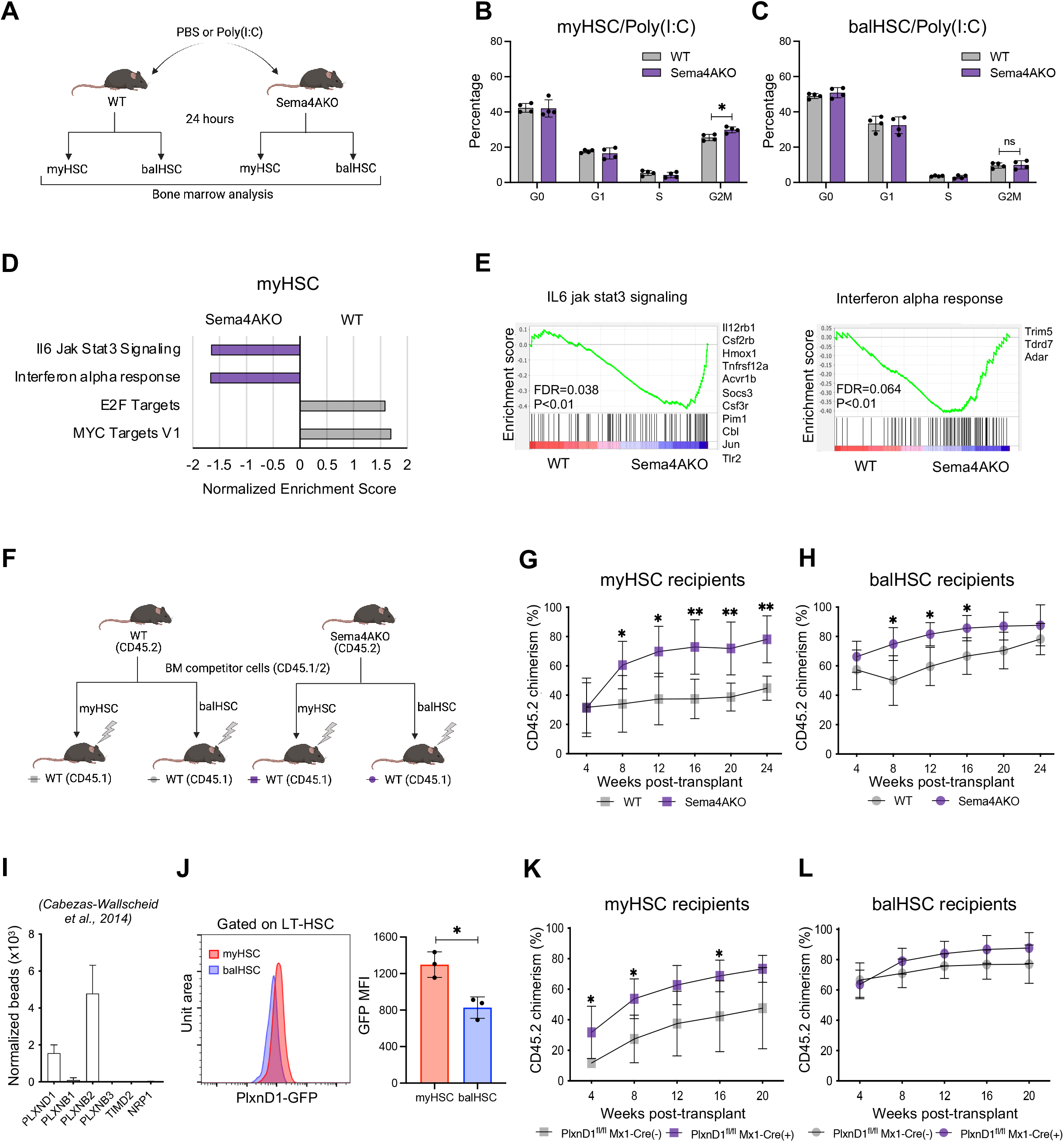
Sema4A/PlxnD1 signaling constrains the response of myeloid-biased HSC to proliferative stress. (A) Experimental schema for the acute inflammatory stress model. (B) Cell cycle analysis of myHSC 24 hours after injection with Poly(I:C) (n=5 mice per genotype). (C) Cell cycle analysis of balHSC 24 hours after injection with Poly(I:C) (n=5 mice per genotype). (D) WT vs Sema4AKO GSEA of myHSC from Poly(I:C) injected mice, FDR<0.01. (E) GSEA plots and top differentially expressed genes for the pathways that were enriched in Sema4AKO myHSC, as shown in (D). (F) Experimental schema for the transplant studies as shown in (G) and (H). (G) Donor chimerism in the recipients of myHSC from young WT/Sema4AKO mice (n=5 mice per donor genotype). (H) Donor chimerism in the recipients of balHSC from young WT/Sema4AKO mice (n=4-5 mice per donor genotype). (I) Published HSC gene expression data (Cabezas-Wallscheid et al., 2014) showing expression of known Sema4A receptors. (J) Representative histograms (left panel) and quantification of mean fluorescence intensity of GFP expression (right panel) in myHSC (pink) and balHSC (blue) from PlxnD1-GFP reporter mice (n=3 mice). (K) Donor chimerism in the recipients of myHSC from PlxnD1^fl/fl^ Mx1-Cre(+)and PlxnD1^fl/fl^ Mx1-Cre (−) mice (n=5 recipient mice per genotype). (L) Donor chimerism in the recipients of balHSC from PlxnD1^fl/fl^ Mx1-Cre(+)and PlxnD1^fl/fl^ Mx1-Cre (−) mice (n=5 recipient mice per genotype). Data are presented as mean ± SD *p < 0.05; **p < 0.01; ***p < 0.001 by unpaired T-test.

Next, we asked if Sema4A deletion differentially impacts long-term reconstitution capacity of the two HSC subsets. To this end, we isolated myHSC and balHSC from WT and Sema4AKO (CD45.2) donors and transplanted equal number of cells from each subset into lethally irradiated WT (CD45.1) recipients (Fig. 2F). As shown in Fig. 2G, Sema4AKO myHSC displayed a significantly higher level of post-transplant reconstitution as compared to WT myHSC controls, with the difference increasing over time (see Fig. S2G for gating strategy used in chimerism analysis). In contrast, this trend was considerably weaker in the recipients of WT/Sema4AKO balHSC and no longer detectable 24 weeks post-transplant (Fig. 2H). In addition, Sema4AKO myHSC (but not balHSC) graft displayed excessive lymphoid skewing (Fig. S2H and S2I). These data provide further support for the myHSC-specific action of Sema4A, as evidenced by enhanced output and impaired differentiation of transplanted Sema4AKO myHSC.

In order to establish a cellular mechanism for this effect, we sought to identify a functional receptor for Sema4A on myHSC. Analysis of published HSC gene expression datasets (Cabezas-Wallscheid et al., 2017; Cabezas-Wallscheid et al., 2014) revealed that amongst known receptors for Sema4A, Plexin-B2 *(PlxnB2*) and Plexin-D1 *(PlxnD1)* had the highest expression level in HSC (Fig. 2I). However, PlxnB2 has been described as a receptor for Angiogenin (Yu et al., 2017) which has no effect on myeloid differentiation (Goncalves *et al.*, 2016). We therefore considered PlxnD1 the most likely candidate. Interestingly, the ability of Sema4A/PlxnD1 signaling to constrain stress-induced proliferation (as it would be for myHSC) has been already shown for the endothelial cells (Toyofuku et al., 2007). Furthermore, our analysis of PlxnD1-GFP reporter mice (Gong et al., 2003) revealed a significantly higher level of PlxnD1 expression in myHSC as compared to balHSC (Fig. 2J), which was consistent with a predominant functional effect of Sema4A. Of note, a fraction of CD34^+^CD90^+^ human HSC also expressed PlxnD1 (Fig. S2J).

Global deletion of PlxnD1 in mice is embryonic lethal due to structural cardiac and vascular defects (Serini et al., 2003), thus precluding functional analysis of adult HSC from these animals. We therefore conditionally deleted PlxnD1 by crossing PlxnD1 “floxed” (Zhang et al., 2009) mice with the Mx1-Cre strain (Kuhn et al., 1995). We confirmed a complete excision of PlxnD1 by PCR and Q-PCR analysis of sorted LKS cells after Poly (I:C) induction (Fig. S2K and S2L). Baseline analysis of PlxnD1^fl/fl^ Mx1-Cre(+) and PlxnD1^fl/fl^ Mx1-Cre(−) mice revealed no significant differences in blood counts, HSC cell cycle and HSPC and mature cell frequency, except for a slight increase in MPP2 and lin^−^Sca-1^−^c-Kit^+^ myeloid progenitors, suggesting myeloid bias (Fig. S2M and S2N, Suppl. Table 1). However, competitive transplantation of myHSC and balHSC from of PlxnD1^fl/fl^ Mx1-Cre(+) and PlxnD1^fl/fl^ Mx1-Cre(−) mice revealed significantly higher reconstitution by PlxnD1-deficient myHSC while their balHSC counterparts engrafted normally, thus recapitulating the phenotype of transplanted myHSC and balHSC from Sema4AKO mice (Fig. 2K-L and S2O-Q). In sum, our results are consistent with a previously unrecognized role of PlxnD1 as a functional receptor for Sema4A on myHSC.

### Sema4A prevents excessive myHSC expansion and functional loss with age

Our observation that Sema4A loss enhances myHSC responsiveness to proliferative challenges, such as acute inflammation and transplantation, prompted us to investigate whether this will lead to impaired myHSC function upon chronic inflammatory stimulation, as occurs during aging (Kovtonyuk et al., 2016). Analysis of peripheral blood in aged Sema4AKO mice revealed progressive anemia, thrombocytosis and neutrophilia (Fig. 3A). In order to rule out systemic inflammation as a cause of the above abnormalities, we examined the plasma levels of 36 proinflammatory cytokines in aged WT and Sema4AKO mice (including thrombopoietin and G-CSF) but detected no significant differences (Suppl. Table 2).

**Figure 3.**
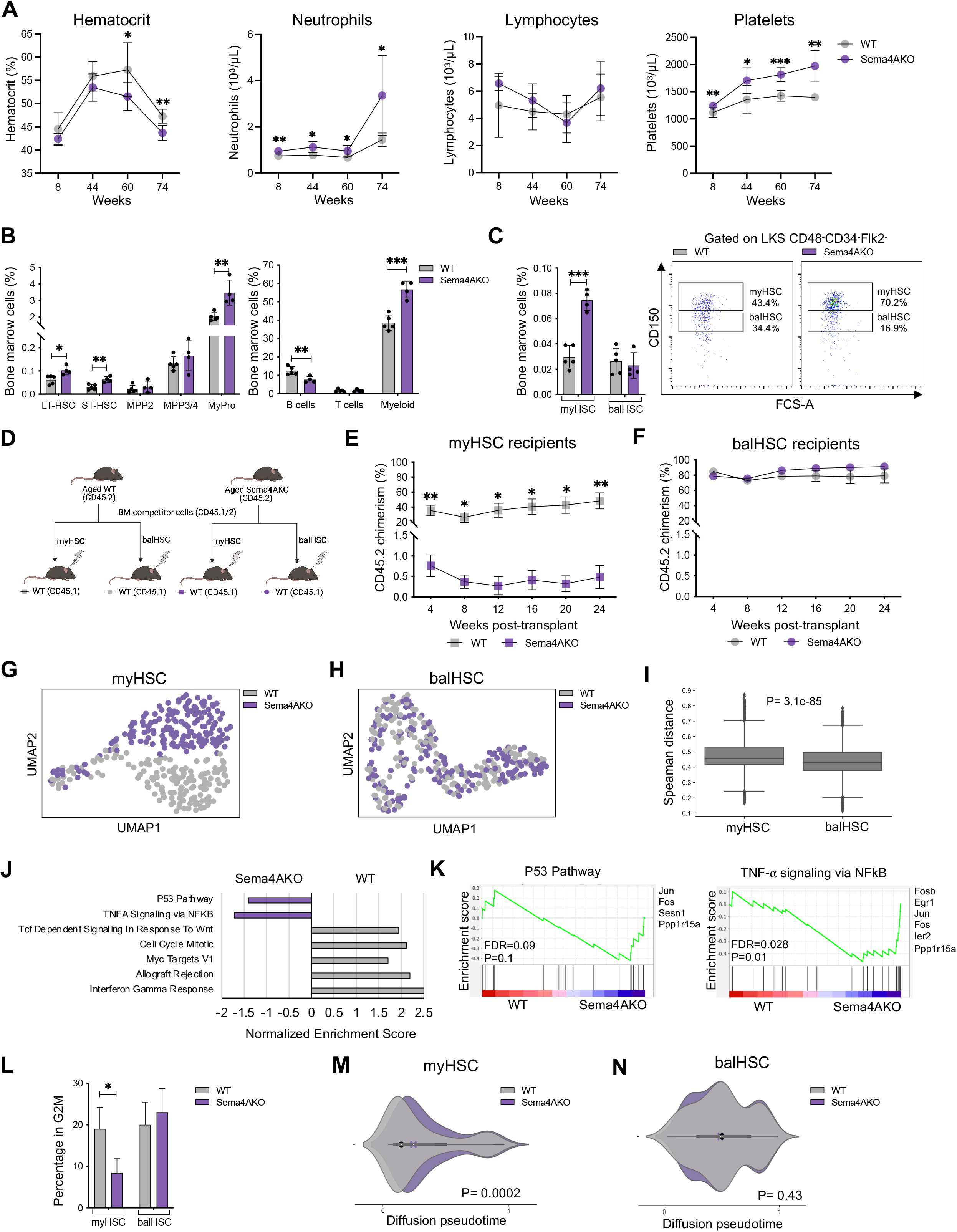
Sema4A prevents excessive myHSC expansion and functional loss with age. (A) Serial peripheral blood counts of WT and Sema4AKO mice during aging (n=4-9 mice per genotype, age range 8-74 weeks). (B) Immunophenotypic analysis of the bone marrow from aged (74-weeks old) WT/Sema4AKO mice (n=4-5 mice per genotype). (C) Frequency of myHSC and balHSC in aged (74-weeks old) WT/Sema4AKO mice (representative flow cytometry plots shown on the right) (n=4-5 mice per genotype). (D) Experimental schema for competitive myHSC/balHSC transplantation experiments shown in and (F). (E) Donor chimerism in the recipients of myHSC from aged WT/Sema4AKO mice (n=3-5 per donor genotype). (F) Donor chimerism in the recipients of balHSC from aged WT/Sema4AKO mice (n=4-5 per donor genotype). (G) UMAP representation of 162 myHSC from aged WT and Sema4AKO mice (n=2 mice per genotype). (H) UMAP representation of 165 balHSC cells from WT and Sema4AKO mice (n=2 mice per genotype). (I) Distribution of pairwise Spearman’s correlation distances between aged WT and Sema4AKO myHSC (left) and balHSC (right). The two distributions are statistically significantly different according to a Wilcoxon rank-sum test (p-value = 3.1e-85). (J) GSEA of myHSC from aged WT/Sema4AKO mice, FDR<0.01. (K) GSEA plots and top differentially expressed genes for pathways enriched in Sema4AKO myHSC. (L) Percentage of myHSC and balHSC cells from aged WT/Sema4AKO in the G2M phase of the cell cycle as estimated from their transcriptome using *Cyclone*. (M, N) Distributions of diffusion pseudotime values of myHSC (panel M) and balHSC (panel N) from aged WT/Sema4AKO. The P-values shown at the bottom were computed with a Wilcoxon-rank sum test. Data are presented as mean ± SD *p < 0.05; **p < 0.01; ***p < 0.001 by unpaired T-test.

Immunophenotypic analysis of the bone marrow in aged Sema4AKO mice demonstrated a higher number of primitive hematopoietic cells and marked myeloid expansion, as evidenced by increased frequency and absolute number of myeloid progenitors and mature myeloid cells (Fig. 3B and S3A). Critically, we observed a marked (~2.5-fold) increase in the absolute number of myHSC while the number of balHSC was comparable with that of aged-matched WT controls (Fig. 3C and S3B).

The amplified aged Sema4AKO myHSC population may represent either expanded *bona fide* myHSC or a more differentiated progeny which retained immunophenotypic features of myHSC but lost long-term regenerative potential following expansion (Bernitz et al., 2016). To distinguish between these two possibilities, we competitively transplanted equal numbers of myHSC from aged Sema4AKO and WT animals into lethally irradiated WT recipients (Fig. 3D).

Strikingly, aged Sema4AKO myHSC, while still displaying myeloid bias, generated a markedly lower level of donor chimerism in peripheral blood (range 0-1.43% vs 9.93-77.5% in aged WT myHSC controls) (Fig. 3E and S3C) and failed to produce a detectable long-term graft in the bone marrow (Fig. S3E). In contrast, balHSC from both WT and Sema4AKO mice gave rise to comparable levels of peripheral blood and bone marrow donor chimerism (Fig. 3F and S3D). These data demonstrate that during aging, Sema4A absence leads to profound functional attrition of phenotypic myHSC but is inconsequential for balHSC.

Aiming to understand the molecular events which are responsible for the myHSC-specific effect of Sema4A, we performed single cell RNA-Seq analysis of myHSC and balHSC from aged WT and Sema4AKO mice using the Smart-Seq2 protocol (Picelli et al., 2014). We sorted 192 cells per group (768 total), of which 642 were selected for analysis following quality control (see Methods). As expected, WT myHSC had a higher expression of *Slamf1*, self-renewal/low-output-associated genes (*CD74*, *Ly6a*, *vWF*, *Procr*) and lower expression of cell cycle-related genes (*Cdk6* and *Mki67*) as compared to WT balHSC (Fig. S3F) (Becker-Herman et al., 2021; Kent et al., 2009; Kent et al., 2008; Laurenti et al., 2015; Morcos et al., 2017; Rodriguez-Fraticelli et al., 2020).

A transcriptome-wide analysis revealed that within the myHSC fraction, WT and Sema4AKO cells formed distinct, minimally overlapping clusters (Fig. 3G) while balHSC of both genotypes merged together, indicating that the absence of Sema4A induces transcriptional changes predominantly in myHSC (Fig. 3H). We quantified this HSC subset-specific difference by detecting a greater correlation-based distance between WT/ Sema4AKO myHSC compared to WT/ Sema4AKO balHSC (Fig. 3I); Wilcoxon Rank-Sum test, p-value 3.1e-85, see Methods for further details).

Next, we examined the transcriptional features of the above aged HSC subsets in more detail. Consistent with the results of the clustering analysis, the number of genes which were differentially expressed in aged Sema4AKO vs WT myHSC was much greater (431) compared to aged Sema4AKO vs WT balHSC (30). In the aged Sema4AKO myHSC signature, we noted markedly reduced expression of genes that normally constrain HSC pool and promote HSC self-renewal (*CD74, vWF, Ly6a, Mllt3*)(Becker-Herman *et al.*, 2021; Calvanese et al., 2019; Kent *et al.*, 2009; Kent *et al.*, 2008), consistent with their excessive expansion and loss of stemness (Fig. S3G). Moreover, GSEA demonstrated a significant enrichment for the terms “p53 pathways” (top genes: *Jun, Fos, Sesn1*) and “TNF-alpha/NFkB signaling” (top genes: *Fosb, Egr1, Jun*) and as well as a recently defined “core aging HSC signature” (Svendsen et al, 2021) (Fig. 3J, 3K, S3H, Suppl. File).

While “TNF-alpha/NFkB signaling” was also enriched in aged Sema4AKO balHSC (Fig. S3I), no enrichment for “p53 pathway” was observed, and enrichment for the “core aging HSC signature” was much weaker (FDR=0.09, P=0.047 for balHSC vs FDR=0.002, P=0.001 for myHSC, (Fig.S3H and data not shown). These findings suggest that in the absence of Sema4A, aged myHSC sustain a greater degree of stress- and inflammation-induced damage (Walter *et al.*, 2015). Consistent with this notion, *in silico* cell cycle analysis (Scialdone et al., 2015) revealed reduced cycling in aged Sema4AKO myHSC but not in balHSC (Fig. 3L). While loss of proliferative capacity in myHSC occurs during normal aging (Montecino-Rodriguez et al., 2019), it was more prominent in aged Sema4AKO myHSC. In conjunction with other phenotypic (expansion and functional loss) and molecular (aging HSC signature) features of normal aging which were exaggerated in aged Sema4AKO myHSC, this suggests that the absence of Sema4A leads to premature myHSC aging.

Given that amplification of aged Sema4AKO phenotypic myHSC was accompanied by a marked expansion of downstream myeloid progeny, we wondered if accelerated differentiation was another factor which would explain their functional loss. We addressed this question using diffusion pseudotime (DPT) analysis (Haghverdi et al., 2016), which can quantify the differentiation state of each cell going from naive (corresponding to HSC) to more mature (multipotent progenitors, MPP). To this end, we first generated 10x Genomics single cell RNA-Seq profiles of lin^−^c-Kit^+^ HSPC from 74-weeks old WT animals, i.e. age-matched with WT/Sema4AKO animals for the Smart-Seq2 single-cell RNA-Seq experiment described above. In this 10x dataset, by mapping expression of previously described markers (Nestorowa et al., 2016) we identified the clusters that correspond to HSC (Cluster 2; *Ly6a, Procr, Hlf*) and MPP (Cluster 0; *Cd34, Cebpa, Ctsg*) (see Methods and Fig. S3J-L). Next, we utilized the transcriptomes of cells within these clusters to estimate a differentiation trajectory (Fig. S3M, see Methods for further details), in which higher DPT values correspond to more mature cells.

As expected, analysis of known self-renewing marker genes revealed downregulation of vWF, *Mpl*, *Fdg5*, C*tnnal1*, *Procr*, and upregulation of *Ctsg* and *Cbpa* as cells progressed from HSC to MPP (Fig. S3N).

We then estimated a DPT value along this trajectory for myHSC and balHSC from aged WT and Sema4AKO mice which were profiled in the Smart-Seq2 experiment described above. Consistent with previously reported myHSC/balHSC hierarchy, the DPT values of WT aged myHSC were lower than WT balHSC, indicating that myHSC are more primitive than balHSC (Fig. S3O), (Carrelha et al., 2018; Morita *et al.*, 2010). Importantly, the comparison between aged WT myHSC and aged Sema4AKO myHSC revealed that the DPT values for aged Sema4AKO myHSC were higher, suggesting that they became more differentiated (p-value = 0.0002, Wilcoxon rank-sum test, Fig. 3M). Conversely, no significant difference was found between the DPT distributions of aged WT and aged Sema4AKO balHSC (Fig.3N). Thus, our DPT analysis demonstrates that the absence of Sema4A during aging leads to premature, myHSC-specific activation of the differentiation transcriptional program.

In sum, our immunophenotypic, functional and transcriptional analysis identified Sema4A as a critical regulator of myHSC self-renewal and differentiation. The profound loss of regenerative capacity in aged Sema4AKO myHSC, as observed in the transplant experiments, likely represents a cumulative effect of partially overlapping cellular defects, such as inflammatory injury, premature aging and accelerated differentiation.

### Sema4A from the bone marrow niche restrains stress-induced myHSC proliferation and maintains self-renewal

Since Sema4A is expressed in both non-hematopoietic and hematopoietic cells, including HSC(Baccin *et al.*, 2020; Cabezas-Wallscheid *et al.*, 2017; Cabezas-Wallscheid *et al.*, 2014), we asked which cellular source of Sema4A was functionally indispensable for myHSC function. First, we investigated the role of HSC-derived Sema4A by employing conditional deletion with Mx1-Cre. We confirmed Cre-induced recombination of the “floxed” Sema4A allele in HSPC by PCR and Q-PCR analysis (Fig. S4A and S4B). Analysis of Sema4A^fl/fl^ Mx-1Cre (+) animals at the steady-state revealed no difference in peripheral blood counts, bone marrow cellularity, cell cycle and frequency of HSPC and mature cells, as compared to Sema4A^fl/fl^ Mx1-Cre(−) controls (Suppl. Table 1, Fig. S4C and S4D). In competitive transplantation assay, no difference in long-term reconstitution capacity of myHSC and balHSC from Sema4A^fl/fl^ Mx1-Cre(+) and Sema4A^fl/fl^ Mx1-Cre(−) mice was observed (Fig. 4A-B and Fig. S4E-G). These data indicate that hematopoietic-derived Sema4A is dispensable for myHSC and balHSC function.

**Figure 4.**
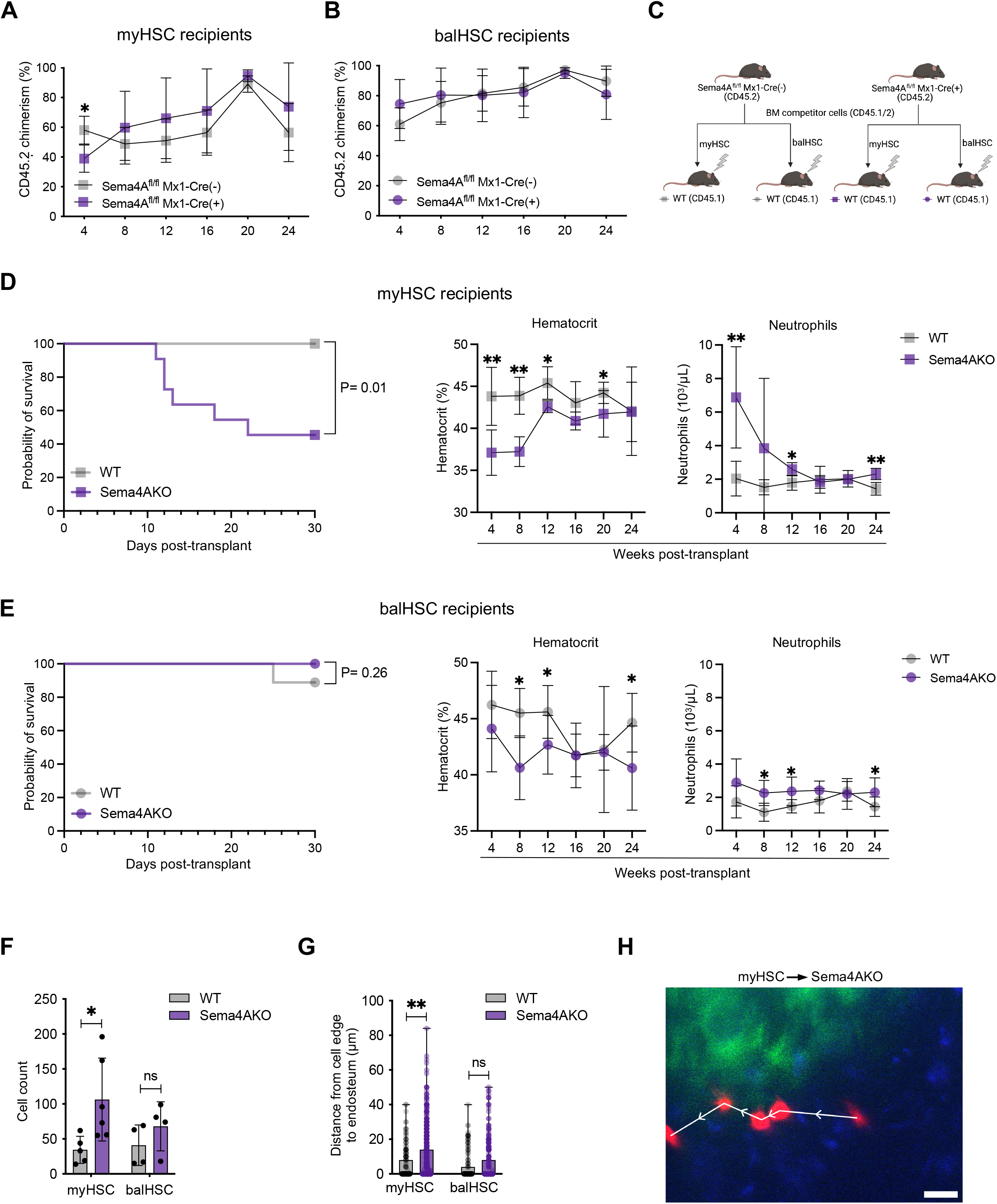
Sema4A from the bone marrow niche restrains stress-induced myHSC proliferation and maintains self-renewal. (A) Donor chimerism in the recipients of myHSC from Sema4A^fl/fl^ Cre(+) and Sema4A^fl/fl^ Cre(−) mice (n=4-5 mice per donor genotype). (B) Donor chimerism in the recipients of balHSC from Sema4A^fl/fl^ Cre(+) and Sema4A^fl/fl^ Cre(−) mice (n=4-5 mice per donor genotype). (C) Experimental schema for non-competitive transplant experiments shown in (D) and (E). (D) Survival curve (left panel) and hematocrit/neutrophil count (right panel) in WT/Sema4AKO recipients of myHSC (data are the summary of 6 independent experiments involving a total of 9-11 recipients per genotype). (E) Survival curve and peripheral blood counts in WT/Sema4AKO recipients of balHSC (data are the summary of 6 independent experiments involving a total of 9-11 recipients per genotype). (F) Average number of cells per mouse ~15-20 hours after transplantation of WT myHSC or balHSC into WT/Sema4AKO recipients, as assessed by two-photon intravital imaging of the calvarial bone marrow (Data are the summary of 6 independent experiments involving a total of 4-6 recipients per genotype). (G) Quantification of 3D distances between individual transplanted cells and the nearest endosteal surface (n = 171, 628, 161, and 266 total cells for WT myHSC, Sema4AKO myHSC, WT balHSC, and Sema4AKO balHSC, respectively; Data are the summary of 6 independent experiments involving a total of 4-6 recipients per group). (H) Representative time-lapse two-photon intravital image of single motile WT myHSC in the calvaria of Sema4AKO recipient. Each cell and white arrow correspond to a different timepoint at increments of 10 mins. The myHSC (DiD, red), bone (SHG, green), and autofluorescence (blue) are shown. Scale Bar ~ 25 μm. Data are presented as mean ± SD *p < 0.05; **p < 0.01; ***p < 0.001 by unpaired T-test.

In order to elucidate the role of bone marrow microenvironment-derived Sema4A, we non-competitively transplanted lethally irradiated WT and Sema4AKO recipients with a radioprotective dose of myHSC and balHSC from WT mice (Fig. 4C). Strikingly, we observed a ~50% post-transplant mortality in Sema4AKO recipients of myHSC (Fig. 4D, left panel). The surviving animals from this group displayed marked anemia and neutrophilia, which resembled blood count abnormalities in aged Sema4AKO mice (Fig. 4D, right panel and Fig. S4H). In contrast, no survival difference was observed in the recipients of balHSC, which showed only mild blood counts changes (Fig. 4E and Fig. S4I). This experiment revealed that Sema4A from the host hematopoietic niche is critical for myHSC self-renewal under stress but plays no significant role in balHSC regeneration.

Among the subsets which make up the bone marrow niche, Sema4A is expressed by endothelial and osteoprogenitor cells (Fig. S1A and Fig. S1B). In order to refine their physiological relevance as cellular sources of Sema4A in the niche, we conditionally deleted Sema4A from each of the two cell types by crossing “floxed” Sema4AKO mice to either VECad-CreERT2 or Osx-Cre animals. Steady-state analysis of Sema4A^fl/fl^ VE-CadCre ERT2(+) and Sema4A^fl/fl^ Osx-Cre (+) mice revealed no difference in peripheral blood counts, bone marrow cellularity, cell cycle and frequency of HSPC and mature cells, as compared to Sema4A^fl/fl^ VE-CadCre ERT2(−) (Sorensen et al., 2009) and Sema4A^fl/wt^ Osx-Cre (+) (Rodda and McMahon, 2006) controls, respectively (Fig. S4J-M and Suppl. Table 1). Transplantation of WT myHSC and balHSC into lethally irradiated donors of the above genotypes showed that both endothelial- and osteoprogenitor-specific deletion of Sema4A only partially recapitulated the effect of a complete Sema4A absence in the host. Specifically, we observed increased mortality in Sema4A^fl/fl^ Osx-Cre (+) recipients of myHSC, but the difference was not statistically significant (Fig. S4N-P). In Sema4A^fl/fl^ VE-CadCre ERT2(+) recipients of myHSC, we detected a slight reduction in hematocrit and no impact on survival (Fig. S4Q-S). Taken together, these studies suggest that a cumulative production by osteoprogenitors, endothelial cells and likely other cellular source(s) may be responsible for the full functional effect of microenvironment-derived Sema4A on myHSC.

Having demonstrated that a complete absence of Sema4A in the host is critical for myHSC engraftment (Fig. 4D), we asked how it may affect early homing, expansion and motility of transplanted myHSC in real time. To this end, we isolated myHSC and balHSC from WT mice and fluorescently labeled them with DiD. We then transferred equal numbers of these cells into lethally irradiated WT and Sema4AKO recipients and performed intravital time-lapse two-photon microscopy of the calvarial bone marrow (Christodoulou et al., 2020). We recorded 3D z-stacks and time-lapse movies 15-20 hours after the cell injection. Single cells and clusters (defined as two or more cells whose cell-to-cell edge are within 15 μm) were detected throughout the calvarial bone marrow in all mice (Fig. S4T). Notably, we observed a ~3 times higher number of transplanted cells in Sema4AKO recipients of myHSC compared to WT controls (mean ~106 cells vs. 34 cells, [p-value = 0.0295]) whereas cell number in WT/Sema4A recipients of balHSC were not significantly different (mean ~68 cells vs. 41 cells, respectively [p-value = 0.2793]) (Fig. 4F and S4U). These results indicate that the absence of Sema4A in the host leads to excessive myHSC expansion but is inconsequential for balHSC. As further evidence for this, we found a similar 3-fold increase in the number of cell clusters in the Sema4AKO recipients of myHSC as compared to WT controls (mean ~21 vs. 7 clusters [p-value = 0.0219] whereas the trend in the balHSC recipients was much weaker (~15 vs. 8 clusters [p-value = 0.2591]) (Fig. S4V).

Next, we analyzed the effect of Sema4A on myHSC and balHSC localization by measuring the 3D distance to the endosteal surface, an established location of post-transplant HSC niche (Lo Celso et al., 2009). Importantly, we observed that in Sema4AKO recipients of myHSC, transplanted cells were found nearly 2x farther from the endosteum compared to WT controls (mean ~8.2 μm vs. 4.9 μm, respectively [p-value = 0.0024]) whereas this difference was smaller and not statistically significant in the Sema4AKO/WT balHSC recipients (mean ~5.9 μm vs. 4.0 μm, respectively [p-value = 0.0674]) (Fig. 4G). These results suggest the in the absence of host Sema4A, myHSC homed away from the niche, whereas localization of balHSC was only marginally altered.

Recent intravital time-lapse microscopy studies revealed that upon proliferative challenge, some HSC within the bone marrow niche become motile (Christodoulou *et al.*, 2020; Upadhaya et al., 2020), indicating that motility may reflect HSC activation state. In order to investigate if motility of transplanted myHSC and balHSC is altered in the absence of Sema4A, we performed time-lapse microscopy for 1.5 hrs. We found that balHSC displayed limited motility (defined as <5 μm movement of the cell centroid over the imaging period) regardless of the host genotype (data not shown). In contrast, a small fraction (~4.7% of total) of myHSC transplanted into Sema4AKO mice exhibited highly motile behavior Sema4A (Fig. 4H, Fig.S4V-X, Suppl. Movie). In sum, our intravital imaging data suggests that host absence of Sema4A leads to myHSC hyperactivation, excessive proliferation and mis-localization, which cumulatively may contribute to the loss of self-renewal and engraftment failure in Sema4AKO recipients of myHSC. Intriguingly, post-transplant behavior of balHSC was relatively unaffected, indicating that the two HSC subsets may have fundamentally different requirements for engraftment, including specific dependence of myHSC on Sema4A.

## DISCUSSION

Our study provides substantial experimental support for the concept that functionally diverse subsets of somatic stem cells are controlled by distinct non cell-autonomous signals. Prior studies have indicated that within the HSC pool, myHSC and balHSC display differential sensitivity to soluble factors, such as TGF-beta, RANTES, CXCL2 and histamine (Challen *et al.*, 2010; Chen et al., 2017; Ergen et al., 2012; Pinho et al., 2018). However, the impact of these molecules on myHSC longevity and interaction with the bone marrow microenvironment has not been investigated in detail.

In the current study, we identify Sema4A as an indispensable and specific regulator of myHSC quiescence and self-renewal. Semaphorins and plexins are large protein families (Alto and Terman, 2017) whose role in regulation of adult stem cell quiescence and self-renewal is not known. We demonstrate that the absence of Sema4A leads to myHSC over-proliferation and hyperactivation following acute inflammatory insult, which correlates with a dramatic loss of regenerative function with age, likely due to the loss of protection from detrimental effects of inflammatory signaling over the animal’s lifetime (Kaschutnig et al., 2015). Notably, WT myHSC are preferentially activated by inflammation (Mann *et al.*, 2018; Matatall *et al.*, 2014; Mitroulis *et al.*, 2018) and become vulnerable to damage, underscoring a physiological need for a dedicated protective signal, such as Sema4A.

Excessive myeloid expansion, as observed in the aged Sema4AKO model, is the cardinal feature of human hematopoietic aging (Pang *et al.*, 2011) and clonal hematopoiesis of indeterminate significance (CHIP) - a common condition which carries a significant risk of progression to myeloid malignancy over time but lacks effective therapeutic intervention (Jaiswal and Ebert, 2019). Our findings raise a possibility that pharmacological augmentation of Sema4A/PlxnD1 signaling may serve as a potential strategy to constrain proliferation of myHSC at the top of expanding myeloid-biased clones, thereby preventing aging-associated HSC dysfunction and reducing the risk of malignant transformation.

Our results underscore the importance of niche-derived signals in life-long maintenance of tissue stem cell hierarchy, which is topped by myHSC in the hematopoietic system. Given that stem cell hierarchies underlie functional organization of other tissues (Altshuler *et al.*, 2021; Farrelly *et al.*, 2021; Hsu *et al.*, 2011; Rompolas *et al.*, 2013; Sachewsky *et al.*, 2019; Scaramozza *et al.*, 2019), the data presented here provide justification for broader efforts to identify subset-specific stem cell regulators, which may lead to development of more precise and effective pro-regenerative therapies.

## LIMITATIONS OF THE STUDY

Recent studies revealed that CD150-expressing phenotypic HSC contain a heterogenous mixture of myeloid-restricted progenitors which vary in their capacity for long-term reconstitution and degree of commitment to myeloid, erythroid and platelet lineage (Yamamoto et al., 2018). We recognize that our experiments are unable to resolve the precise identity of a myeloid-restricted subset(s) which is regulated by Sema4A beyond the CD150^high^ HSC fraction. Future studies, including single cell transplantation experiments, will be required to address this question.

Although our data demonstrates that Sema4A suppresses myHSC proliferation during stress, the downstream mediators of this effect are not known. Therefore, the molecular consequences of Sema4A binding to PlxnD1 (and potentially other Sema4A receptors whose functional relevance has not be ruled out by the current study) will need to be further investigated.

## ACKNOWLEDGMENTS

Work in the Silberstein Laboratory is supported by the NIH/NHLBI (R01 HL148189), Translational Research Program grant through Leukemia and Lymphoma Society, Washington State CARE Fund and New Development Fund from Fred Hutchinson Cancer Research Center. The work in the Göttgens laboratory was funded in part by the Wellcome Trust [203151/Z/16/Z] and the UKRI Medical Research Council [MC_PC_17230]. For the purpose of open access, the author has applied a CC BY public copyright license to any Author Accepted Manuscript version arising from this submission. The work in the Maes laboratory is supported by grants from the Research Foundation Flanders (FWO) and KU Leuven (C1). The schematics were created with BioRender.com.

We would like to thank Elena Nefyodova (KU Leuven) for expert technical help, Michael Li, Zoe Derauf, Ben Janoschek and Kora Krumm for mouse colony support, and the staff of Flow Cytometry, Genomics and Comparative Medicine cores facilities at Fred Hutchinson Cancer Research Center. We are also grateful to Peter Klein, University of Pennsylvania, and Adam Wilkinson, University of Oxford, for helpful feedback and advice.

## AUTHOR CONTRIBUTIONS

DT and SZ designed and performed experiments, analyzed and interpreted data, and wrote the manuscript. EM performed computational analyses of sequencing data and contributed to data interpretation. EIC performed experiments, computational analyses of sequencing data and contributed to experimental design and data interpretation. AS data supervised all the computational analyses of sequencing data and contributed to data interpretation. AP supported mouse breeding and performed experiments. NT, CB and DG performed intravital imaging experiments under the supervision of JAS. SR, HPK designed and performed human HSC experiments. NKW and SJK performed single cell RNA-Seq experiments under the supervision of BG. CN, MM, CM and PK contributed data. SR and NC performed experiments with aged mice. TW and AK contributed reagents. EMP and DTS contributed to experimental design and data discussion. L.S. conceived and supervised the project, performed experiments, analyzed data and wrote the manuscript.

