## Supplemental Information for "Myeloid-biased HSC require Semaphorin 4A from the bone marrow niche for self-renewal under stress and life-long persistence"

### RESOURCE AVAILABILITY

#### Lead contact

#### Materials availability

This study did not generate new unique reagents.

#### Data and code availability

All data were analyzed with standard programs and packages, as detailed in Methods. Sequencing data from this study are being submitted to Array Express and expected to be available shortly.

### EXPERIMENTAL MODEL AND SUBJECT DETAILS

#### Animals

All animal experiments were approved by the Institutional Animal Care and Use Committee at Fred Hutchinson Cancer Research Center. All intravital imaging experiments were performed according to procedures approved by the Institutional Animal Care and Use Committee at the University of California, Merced. Wild-type C57Bl/6J, B6SJL, Ubiquitin-GFP, Mx1-Cre, and Osx-Cre mice were obtained from Jackson laboratory. Sema4AKO mice were obtained from Dr. A Kumanogoh, University of Osaka (Kumanogoh et al., 2005). VE-CadCreERT2 mice were obtained from Max-Planck Institute. PlxnD1 conditional KO mice and PlxnD1-GFP were obtained from Dr Chenghua Gu, Harvard University. Sema4A conditional KO mice were obtained from Dr T Worzfeld, University of Marburg. Generation of mice carrying a targeted allele of Sema4A has been described previously (Xia et al., 2015). These mice were crossed with Flp mice to obtain a floxed Sema4A allele.

To induce conditional deletion via CreERT2, 6-8-week-old mice were put on Tamoxifen-containing chow (250mg/kg, Envigo) for 4 weeks followed by normal chow for 4 weeks prior to analysis. To induce conditional deletion via Mx1-Cre, 6-8-week-old mice were interperitoneally (I.P.) injected with 100 µg/mouse high molecular weight polyinosine-polycytidylic acid (Poly(I:C) HMW, InvivoGen) once every other day for a total of five injections and analyzed 4 weeks after the last injection.

For all experiments, both male and female animals were used. Each cohort of animals was matched for age and sex. Young mice ranged from 8-12 weeks old and aged mice ranged from 74-80 weeks old. For the aging experiments, both male and female cohorts were generated but only the female cohort was analyzed in detail. Both WT and Sema4AKO mice were aged in the same environment.

#### PCR genotyping and excision validation for conditional alleles

Genomic DNA was extracted from peripheral blood samples using the DNeasy Blood & Tissue Kit (Qiagen) according to the manufacturer’s instruction or using a fast extraction method involving incubation in ACK Lysing Buffer (Gibco) followed by resuspension in 50mM NaOH, incubation at 95C, and neutralization with 1M tris buffer (pH 7). Genotyping PCR primer sequences are listed in the reagents table.

To confirm Mx1-Cre-mediated excision of Sema4A or PlxnD1 “floxed” allele, up to 50,000 LKS (lin$^-$c-Kit$^+$Sca-1$^+$) cells from animals of desired genotypes were sorted into 350µL RLT Plus Buffer (Qiagen). DNA and RNA were extracted with AllPrep DNA/RNA Micro Kit (Qiagen) according to the manufacturer’s instructions. Primers listed in the reagent table were used to detect the excised and non-excised alleles by genomic DNA PCR. In addition, RNA was reverse transcribed using SuperScript IV VILO Master Mix (Invitrogen) and expression of Sema4A exon 1 and PlxnD1 exon 1 relative to GAPDH was determined using qPCR primers listed in the reagent table.

**Flow cytometry and cell sorting**

Whole bone-marrow mononuclear cells (BMMNC) were collected by crushing tibias, femurs, and pelvis in Ca2+/Mg2+-free phosphate-buffered saline (D-PBS) supplemented with 2% fetal bovine serum (FBS, Fisher Scientific). For mature cells analysis, BMMNC were stained with fluorochrome-conjugated Mac1 (Invitrogen), Gr1 (BioLegend), B220 (BioLegend), and CD3ε (BioLegend) antibodies for 30 minutes. For hematopoietic stem and progenitor cell (HSPC) analysis, BMMNC were stained with fluorescently conjugated Sca-1 (BD Biosciences), c-Kit (BD Biosciences), Lineage cocktail (BD Biosciences), CD48 (BD Biosciences), CD34 (BD Biosciences), Flk2 (BD Biosciences), and CD150 (BioLegend) antibodies for 90 minutes. For cell sorting, BMMNC were lineage depleted by staining with CD3ε, CD11b, B220, TER-119, Gr-1, CD4 and CD8α biotin-conjugated antibodies (all from BD Biosciences) followed by application of streptavidin microbeads (Miltenyi Biotec) and depletion using magnetic separation columns (Miltenyi Biotec). Lineage-negative fraction was stained with conjugated monoclonal antibodies for HSPC markers as described above, except fluorescently conjugated streptavidin replaced fluorescently conjugated lineage cocktail. Samples were analyzed on BD FACSymphony A5 (BD Biosciences) or sorted on BD Aria II (BD Biosciences). All flow cytometry data were analyzed using FlowJo software. Graphs were made using Prism 9 (GraphPad) software or Microsoft Excel.

**Cell Cycle analysis/EdU incorporation**

For cell cycle analysis, we used our recently developed protocol (Galvin et al., 2019). BMMNC were stained with conjugated monoclonal antibodies for HSPC markers, as described above. The cells were fixed and permeabilized using Cytofix/Cytoperm™ Fixation/Permeabilization kit (BD Biosciences) according to the manufacturer’s instruction. Cells were then stained with fluorescently conjugated Ki67 antibody (1:20, BD Biosciences) for 45 minutes followed by DAPI (1:300, Invitrogen) for 10 minutes. Samples were analyzed at 2500-3000 threshold rate. For 5-ethynyl-2’-deoxyuridine (EdU) incorporation, mice were injected with 1mg/mouse EdU 3 hours prior to analysis. Following lineage depletion, the lineage-negative fraction was stained with conjugated monoclonal antibodies for HSPC markers, as described above. Cells were then fixed, permeabilized, and stained for EdU using Click-iT™ Plus EdU Alexa Fluor™ 488 Flow Cytometry Assay Kit (Invitrogen) according to the manufacturer’s instruction. All samples were analyzed using BD FACSymphony A5 (BD Biosciences).

**Transplantation experiments**

For all transplantation experiments, recipients were lethally irradiated 24 hours prior to transplant at 1200 cGy (a split dose of 600 cGy + 600 cGy with a 3 hr interval between doses) and maintained on Baytril-containing water for the first 4 weeks.

For competitive transplantation experiments involving young mice, 200 myHSC (lin$^-$c-Kit$^+$Sca-1$^+$CD48$^-$CD34$^-$Flk2$^-$CD150$^{high}$) or 200 balHSC (lin$^-$c-Kit$^+$Sca-1$^+$CD48$^-$CD34$^-$Flk2$^-$CD150$^{low}$) from CD45.2 C57Bl/6J, Sema4AKO, PlxnD1$^{fl/fl}$ Mx1-Cre and Sema4A$^{fl/fl}$ Mx1-Cre mouse models were co-injected with 200K CD45.1/2 competitor cells. For competitive transplantation experiments using HSC from aged mice, 2000 myHSC or 2000 balHSC CD45.2 cells were transplanted with 200K CD45.1/2 competitor cells. Peripheral blood donor chimerism was monitored at 4-week intervals. Flow cytometry gating strategies for the identification of myHSC and balHSC are shown in Supplementary Figure 1.

For the non-competitive transplant experiments, the animals were lethally irradiated (as described above) and injected with 1100-1600 WT (CD45.1) myHSC or balHSC. Engraftment kinetics were monitored by blood counts and flow cytometry every 4 weeks.

**Post-transplant chimerism analysis**

Peripheral blood was collected from each recipient via retro-orbital sinus, and complete blood count (CBC) was performed immediately after blood collection using Element HT5 (HESKA). Red blood cells were lysed in ACK Lysing Buffer (Gibco) and then stained with fluorescently conjugated CD45.1 (BioLegend), CD45.2 (BioLegend), Mac1 (Invitrogen), Gr1 (BioLegend), B220 (BioLegend), and CD3 (BioLegend) antibodies. Samples were analyzed using BD FACSymphony A5 (BD Biosciences).

**Induction of inflammatory response**

High molecular weight polyinosine-polycytidylic acid (Poly(I:C) HMW, InvivoGen) was prepared according to the manufacturer's instructions. For Poly(I:C)-induced inflammatory studies, mice were interperitoneally injected with 10mg/Kg Poly(I:C) (Jalbert and Pietras, 2018) and analyzed 24 hours after injection. Ultrasound was used to guide intraperitoneal injections.

**Human CFSE assay**
Human GCSF-mobilized $CD34^+$ cells were obtained from Co-Operative Center for Excellence in Hematology at Fred Hutchinson Cancer Research Center. The cells were stained with CellTrace™ CFSE (Invitrogen) according to the manufacturer's recommendation. Briefly, $CD34^+$ cells at $10^6$/ml were incubated with CellTrace™ CFSE stock solution for 20 minutes at 37C. $CD34^+$ cells were seeded in culture media (StemSpan™ SFEM II, Stem Cell Technologies) supplemented with 1% Penicillin-Streptomycin (Gibco), as well as SCF, TPO, and FLT-3 at 100ug/mL (all three PeproTech). Cells were seeded at a density of $10^6$/ml. h-Sema4A-Fc or hIgG1 protein (0-10ug/ml, R&D Systems) was added to the plated cells. Samples were analyzed using BD FACSymphony A5 (BD Biosciences) 48-72hr after seeding the cells.

**Mouse colony-forming unit assay**

25000 Whole bone-marrow mononuclear cells from young WT C57Bl/6J mice were plated with mSema4A-Fc-hIgG1 (Fred Hutchinson Cancer Research Center Molecular Design and Therapeutics) or hIgG1 (Fred Hutchinson Cancer Research Center Molecular Design and Therapeutics) at the indicated concentrations in MethoCult GF M3434 (Stem Cell Technologies) and cultured at 5% $CO_2$ and 37 C. Colonies were manually counted on day 14.

**Mouse LKS cell culture**

1000 LKS cells from Ubiquitin-GFP mice (The Jackson Laboratory) were sorted into 384-well plates (Corning) and cultured in serum-free S-clone SF-O3 medium (Iwai North America Inc.) supplemented with 1% bovine serum albumin (New England BioLabs), mouse SCF (50 ng/mL; PeproTech),TPO(50 ng/mL; PeproTech) and hSema4A-Fc-hIgG1 (LSBio) or hIgG1 (LSBio) at the indicated concentrations. Each well was imaged on Olympus IX81 fluorescent microscope, and the number of $GFP^+$ cells was quantified using ImageJ software.

**Human Sema4A fluorescence in situ hybridization**

Discarded bone tissue from hip replacement surgery was obtained from the Department of Orthopedic Surgery, University of Washington, fixed in formalin, decalcified in EDTA, frozen and sectioned. Human Sema4A probe was obtained from Advanced Cell Technologies. FISH was performed according to the manufacturer's protocol.

**Cytokine array**

ELISA was performed on plasma samples using Cytokine & Chemokine 36-Plex Mouse ProcartaPlex™ Panel 1A, TGF beta 1 Mouse ProcartaPlex™ Simplex Kit, ProcartaPlex Mouse Basic Kit, and Thrombopoietin (TPO/THPO) Mouse ELISA Kit (all from Invitrogen).

**Intravital imaging**

MyHSC and balHSC were isolated by flow sorting, as described above. Cells were stained with 10 µM DiD (Invitrogen) in D-PBS supplemented with 2% FBS for 20 minutes at 37°C. ~1500 DiD-stained myHSC and balHSC were suspended in ~100 µl of $Ca^{2+}$/$Mg^{2+}$-free phosphate-buffered saline (D-PBS) and adoptively transferred via retro-orbital injection into anesthetized young WT and Sema4AKO mice. One day before transplantation, recipient mice were lethally irradiated (1200 cGy) using an x-ray irradiator (X-RAD 320, Precision) with a split dose of 600 cGy + 600 cGy with a 3 hr interval between doses.

For intravital imaging, mice were prepared as previously described (Christodoulou et al., 2020). Briefly, 14-15 hours after transplantation, the mice were anaesthetized with an induction dose of 3–4% isoflurane and a maintenance dose of 1.5–2% isoflurane. Mice were deemed anaesthetized by the toe pinch method. The hair on the calvarium was removed with a mechanical trimmer and the skin was cleaned with alcohol. The mice were mounted in a custom-designed heated mouse holder (for z-stack or time-lapse imaging). Next, a calvarial skin flap was created with a U-shaped incision to reveal the underlying calvaria as previously described. A drop of D-PBS was applied to the skull as the immersion fluid. The mice were transferred to the stage of a multiphoton/confocal laser-scanning video-rate microscope and an Olympus 25× 1.05 numerical aperture water-dipping objective was used for the imaging. Regions 3 and 4 of the calvarial bone marrow were imaged for ~3 hours per mouse and DiD cells were located and imaged (Christodoulou *et al.*, 2020; Sipkins et al., 2005). Z-stacks were acquired with 2 µm steps and time-lapse images were acquired at 10 min intervals for 90 min in 4-8 fields of view. The excitation wavelength from Insight X3 (Spectra-Physics) and Mai-Tai eHP (Spectra-Physics) lasers were set to 1040 nm for two-photon excitation of DiD (emission collected with a 659-700 nm bandpass filter) and second harmonic generation for bone imaging (emission collected with a 503-538 nm bandpass filter), and 820 nm for two-photon excitation of autofluorescence (emission collected with a 572-608 nm bandpass filter). After imaging was completed, mice were euthanized according to approved animal protocols. The contrast and brightness of images and videos were adjusted for display purposes only.

Intravital images were processed and analyzed using Fiji (ImageJ 1.53k). Representative images of single cells and clusters were created by taking Maximum Intensity Projections (MIPs) of z-stacks and adjusting the image contrast. The number of transplanted cells in the R3/R4 region of the calvaria was quantified and the distance from the transplanted cell's edge to the nearest endosteum surface was determined for each cell. Transplanted cells were classified as single or cluster cells when the nearest cell-to-cell edge distance was >15 µm or < 15 µm, respectively.

Motile transplanted cells were identified as those cells whose centroid moved at least 5 µm over the course of 90 min. Cell motility was determined by taking MIPs of the timelapse z-stacks and aligning them with the "Linear Stack Alignment with SIFT" Fiji plugin. The cell centroid at each timepoint was determined manually and the XY displacement between each timepoint was calculated.

**Bulk RNA Sequencing**

For myHSC/balHSC profiling from poly(I:C)-injected mice, 50 cells from each fraction of interest were sorted into the lysis buffer -10% Triton X-100 (Sigma-Aldrich), SUPERase-In RNase Inhibitor 20U/µl (Ambion) and snap-frozen. cDNA amplification was performed as per Smart-Seq2 protocol. Fragmentation and sample barcoding was performed using the NEBNext® Ultra™ II FS DNA Library

Prep Kit for Illumina (New England Biolabs) according to manufacturer's guidelines. Samples were sequenced on a 200 cycle NovaSeq SP.

**Single-cell RNA sequencing**

*HSPC.* BMMNC were stained with conjugated monoclonal antibodies for HSPC markers as described above. 10,000 $Lin^{-}kit^{+}$ cells from young or aged WT/Sema4AKO mice were sorted in Ca2+/Mg2+-free phosphate-buffered saline (D-PBS) supplemented with 2% bovine serum albumin (New England Biolabs). Following one round of centrifugation, the cells were processed on 10x Genomics platform according to the manufacturer's protocol.

*MyHSC/balHSC.* BMMNC were stained with conjugated monoclonal antibodies for HSPC markers as described above except for CD34 in order to minimize staining time and prevent RNA degradation (CD34 staining requires 90 minutes (Okamoto et al., 2007). MyHSC and balHSC were sorted into 96-well plates with the lysis buffer and stored frozen at -80. cDNA amplification and sequencing were performed as per Smart-Seq2 protocol (Picelli et al., 2013).

**Bulk sequencing analysis**

The quality of the reads including Phred quality score (quality of the identification of nucleobases during sequencing) adaptor contamination, GC content, duplicate levels was assessed using the tool FastQC (Andrews, 2010). Illumina adaptor sequences were trimmed off reads using the tool Flexbar (Dodt et al., 2012). Reads were mapped to the human reference genome hg38 (Ensembl version 85) using STAR (Dobin et al., 2013). SAMtools (Li et al., 2009) was used to sort and index the aligned reads. The multicov function of BEDtools (Quinlan and Hall, 2010) was used to count the read fragments per gene and generate a matrix of reads per gene.

In R, the tool DESeq2 (Love et al., 2014) was used for DGE analysis which provides log2 fold change of genes between two conditions. P values were adjusted for by multiple hypothesis correction with the Benjamini and Hochberg method to produce adjusted p. values (p.adj) (Benjamini et al., 2001). Genes were considered significant with a p.adj < 0.01. For pathway enrichment analysis the full list of differentially expressed genes were ranked by statistical significance and the GSEA tool from the Broad Institute was used to perform enrichment analysis (Subramanian et al., 2005). In particular, annotated gene sets from the following databases were used: Hallmark gene sets (Liberzon et al., 2015), BioCarta, KEGG (Kanehisa and Goto, 2000), Reactome (Croft et al., 2011) and Gene ontology (GO) (Ashburner et al., 2000; Gene Ontology, 2021). Pathways with a false discovery rate (FDR) of < 0.1 were considered significant. Overlapping pathways were identified and consolidated using Cytoscape (Shannon et al., 2003) with the Enrichment Map plugin (Merico et al., 2010).

**Raw data processing and normalization**

We quantified the abundance of the transcripts from 768 cells with salmon (v0.17) (Patro et al., 2017). First, we indexed the mouse transcriptome (GRCm38) in quasi-mapping-based mode with --seqBias and --gcBias flags. Then, we aggregated the transcript level abundances into gene level abundances, which in turn was transformed into a gene-cell count matrix. Next, quality control was performed to filter out cells that satisfy any of the following criteria: 1) less than 4000 genes detected (detection threshold: reads-per-million>10); 2) overall mapping less than 50%; 3) fraction of reads mapping to mitochondrial transcripts larger than 0.02; 4) fraction of reads mapping to ERCC spike-ins higher than 0.01. After quality control, we retained 642 cells for downstream analyses (155 WT myHSC, 157 WT balHSC, 166 KO myHSC, 164 KO balHSC). The data were normalized using 'quickcluster' and 'computeSumFactors' functions from the scran package in R (Lun et al., 2016). Finally, we added a pseudocount of 1 to the count matrix, followed by natural-log transformation.

**Batch integration and visualization**

Since the data were collected from two separate batches, we performed batch integration before visualizing the data. For this purpose, we used the Seurat batch integration workflow (Butler et al., 2018). First, 3000 highly variable genes were selected from each batch. Then, we used the FindIntegrationAnchors function to estimate the anchors to use for the integration with 3000 features (anchor.features =3000) and the first 20 canonical variates (dims =1:20). Finally, the 'IntegrateData' function was used with default parameters to integrate the two batches.
To visualize the cells on a low dimensional space, we first built a k-nearest neighbor graph with 'neighbors' function from scanpy (Wolf et al., 2018) with 10 principal components (PCs) and k=30. Then, the 'tl.umap' function was used to calculate a UMAP representation (Becht et al., 2018) and first two UMAP components were plotted. We applied this procedure separately for myHSC and balHSC.
To verify if myHSC and balHSC are differentially affected by the absence of Sema4A, we calculated the pairwise Spearman's correlation distance (defined as (1-ρ)2, where ρ is the Spearman's correlation coefficient computed on the top 3000 highly variable genes identified with Seurat) between WT and KO cells for myHSCs and balHSC separately. Then, we tested the statistical significance of the difference between the two distributions of pairwise distances by using the Wilcoxon rank-sum test.

#### Cell cycle prediction

We estimated the cell cycle phase of each cell by applying the "pairs" algorithm, described by Scialdone et al. (2015) using the implementation in the scran package. More specifically, we used the function 'cyclone' with a minimum number of pairs 120 (mm.pairs=120). To test the difference in percentage of cells allocated to G2/M phase between WT and KO myHSC, we first merged the number of cells allocated to G1 and S phases for each condition. Finally, we built a contingency table and performed Pearson's chi-square test with 'chi2_contingency' function from the 'scipy' module in python.

#### Diffusion pseudotime analysis

For this analysis, we considered the clusters of HSCs and multipotent progenitors (MPPs) from the 10x WT aged mice (see below), and we integrated them with the Smart-seq2 myHSC dataset using the Seurat integration pipeline as described above. Next, we built a diffusion map to establish a differentiation trajectory. For this purpose, a k-nearest neighbor graph was first built with the first 5 PCs and k=15. A diffusion map was then constructed with the 'tl.diffmap' function from scanpy. We defined a diffusion pseudotime (DPT) by selecting the root cell that had the lowest value of the first diffusion component. Finally, in order to test the difference in the distribution of DPT values for both WT and KO myHSCs, we used the Wilcoxon rank-sum test.

#### Differential gene expression analysis

We found differentially expressed genes between KO and WT with the DEseq2 package (Love *et al.*, 2014) on R. First, we removed genes that were expressed in fewer than 10 cells for each batch, and the counts were rounded to integers. Then, we created a DESeq object from the count matrix with the design ~condition+batch. Fold-changes and p-values were estimated using the DESeq function with default parameters.

#### 10x Genomics library preparation

Libraries from 10,000 Kit+ cells each from WT (n=1) and Sema4A KO (n=2) young and WT (n=2) old mice were prepared using the Single Cell 3′ Reagent Kit v3 (10x Genomics) according to the manufacturer's protocol.

#### Quality control and data normalization

The Cell Ranger 3.1.0 (10x Genomics) analysis pipeline was used to process the 10x single cell RNA-Seq output by aligning reads to mm10-3.0.0 mouse transcriptome (Ensembl). Cell Ranger was also used to aggregate condition replicates prior to processing. For all the analysis steps specified below, we used the SCANPY library (Wolf *et al.*, 2018), unless specified otherwise. As a quality control, first we filtered out cells with fewer than 200 genes detected with >0 reads, and we removed genes present in fewer than 3 cells. We also removed cells with a high percentage of reads mapped to mitochondrial genes (>5%) and a high number of detected genes (>6500). With the tool Scrublet (Wolock et al., 2019), we predicted and removed doublets from the dataset.
After concatenating the data from WT and Sema4AKO mice, we normalized the data for sequencing depth to a target of 1e4 counts per cell. Then, after adding 1 as pseudocount, the count matrix was log-transformed. Finally, the total counts per cell as well as the percentage of reads mapping to mitochondrial genes were regressed out.

**Data clustering and visualization**

We identified highly variable genes with the scanpy function “sc.pp.highly_variable_genes” (parameters: min_mean=0.0125, max_mean=3, min_disp=0.5) and we computed a neighborhood graph (n_neighbors = 8) on the first 40 principal components. To visualize the data, the graph was embedded in 2 dimensions using Uniform Manifold Approximation and Projection (UMAP; see Supplementary Figure 1 (WT and Sema4AKO young) and 3 (WT Old)) (McInnes et al., 2020).

We clustered the neighborhood graph using the Leiden clustering algorithm (Traag et al., 2019) with resolution 1 and 0.3 for young and old mice respectively. This resulted in 28 clusters in the young mice dataset and 17 clusters in the old mice dataset. The WT and Sema4AKO young dataset was subsetted to exclude small, isolated clusters and re-clustered with resolution 0.2. Cluster marker genes were found with a Wilcoxon rank-sum test (see Supplementary Figure 1). Based on known HSC markers (Ly6a, Procr, Hoxb5), we identified the cluster corresponding to HSC in the dataset from WT and Sema4AKO young mice (cluster 1), which was used for the enrichment analysis reported in Figure 1. In the WT old mice dataset, we similarly identified HSC and early multipotent progenitors (MPP) using marker genes and by predicted lineage relationships using graph abstraction (Wolf et al., 2019) (see Supplementary Figure 3), and, based on them, we built a differentiation trajectory for DPT analysis (see above).

**Differential gene expression and enrichment analysis**

We identified the genes differentially expressed between the WT and Sema4AKO cells in the HSC cluster 1 from young mice using a Wilcoxon test. For the pathway enrichment analysis shown in Figure 3, the top 500 upregulated and downregulated genes were ranked by statistical significance and the GSEA tool from the Broad Institute was used to perform enrichment analysis as described for bulk sequencing (Subramanian *et al.*, 2005). An aging signature pathway was also generated using the 220 gene signature identified in Svendsen et al., 2021, Supplementary File 2 - ‘Aging signature - Re-analysis tab’ (Flohr Svendsen et al., 2021).

**Supplementary Movie**

Supplementary Movie 1: Time-lapse movie of motile DiD-labeled myHSC (from Fig. 4H) transplanted into Sema4AKO recipient. Images were recorded every 10 min. from same field of view. The myHSC (DiD, red), bone (SHG, green), and autofluorescence (blue) are shown. Scale bar ~25 μm.

**Supplementary File**

Summary of the RNAseq data (GSEA terms and list of differentially expressed genes).

**Supplementary Table 1**

Baseline blood counts for PlxnD1$^{fl/fl}$ Mx1-Cre, Sema4A$^{fl/fl}$ VE-CadCreERT2, Sema4A Osx-Cre(+) and Sema4A$^{fl/fl}$ Mx1-Cre.

**Supplementary Table 2**

Cytokine array on plasma samples from aged WT and Sema4aKO mice.

### METHOD REFERENCES


Andrews, S. (2010). FastQC: A Quality Control Tool for High Throughput Sequence Data [online]. http://www.bioinformatics.babraham.ac.uk/projects/fastqc/

Ashburner, M., Ball, C.A., Blake, J.A., Botstein, D., Butler, H., Cherry, J.M., Davis, A.P., Dolinski, K., Dwight, S.S., Eppig, J.T., et al. (2000). Gene ontology: tool for the unification of biology. The Gene Ontology Consortium. Nat Genet *25*, 25-29. 10.1038/75556.

Becht, E., McInnes, L., Healy, J., Dutertre, C.A., Kwok, I.W.H., Ng, L.G., Ginhoux, F., and Newell, E.W. (2018). Dimensionality reduction for visualizing single-cell data using UMAP. Nat Biotechnol. 10.1038/nbt.4314.

Benjamini, Y., Drai, D., Elmer, G., Kafkafi, N., and Golani, I. (2001). Controlling the false discovery rate in behavior genetics research. Behav Brain Res *125*, 279-284. 10.1016/s0166-4328(01)00297-2.

Butler, A., Hoffman, P., Smibert, P., Papalexi, E., and Satija, R. (2018). Integrating single-cell transcriptomic data across different conditions, technologies, and species. Nat Biotechnol *36*, 411-420. 10.1038/nbt.4096.

Christodoulou, C., Spencer, J.A., Yeh, S.A., Turcotte, R., Kokkaliaris, K.D., Panero, R., Ramos, A., Guo, G., Seyedhassantehrani, N., Esipova, T.V., et al. (2020). Live-animal imaging of native haematopoietic stem and progenitor cells. Nature *578*, 278-283. 10.1038/s41586-020-1971-z.

Croft, D., O'Kelly, G., Wu, G., Haw, R., Gillespie, M., Matthews, L., Caudy, M., Garapati, P., Gopinath, G., Jassal, B., et al. (2011). Reactome: a database of reactions, pathways and biological processes. Nucleic Acids Res *39*, D691-697. 10.1093/nar/gkq1018.

Dobin, A., Davis, C.A., Schlesinger, F., Drenkow, J., Zaleski, C., Jha, S., Batut, P., Chaisson, M., and Gingeras, T.R. (2013). STAR: ultrafast universal RNA-seq aligner. Bioinformatics *29*, 15-21. 10.1093/bioinformatics/bts635.

Dodt, M., Roehr, J.T., Ahmed, R., and Dieterich, C. (2012). FLEXBAR-Flexible Barcode and Adapter Processing for Next-Generation Sequencing Platforms. Biology (Basel) *1*, 895-905. 10.3390/biology1030895.

Flohr Svendsen, A., Yang, D., Kim, K., Lazare, S., Skinder, N., Zwart, E., Mura-Meszaros, A., Ausema, A., von Eyss, B., de Haan, G., and Bystrykh, L. (2021). A comprehensive transcriptome signature of murine hematopoietic stem cell aging. Blood *138*, 439-451. 10.1182/blood.2020009729.

Galvin, A., Weglarz, M., Folz-Donahue, K., Handley, M., Baum, M., Mazzola, M., Litwa, H., Scadden, D.T., and Silberstein, L. (2019). Cell Cycle Analysis of Hematopoietic Stem and Progenitor Cells by Multicolor Flow Cytometry. Curr Protoc Cytom *87*, e50. 10.1002/cpcy.50.

Gene Ontology, C. (2021). The Gene Ontology resource: enriching a GOld mine. Nucleic Acids Res *49*, D325-D334. 10.1093/nar/gkaa1113.

Jalbert, E., and Pietras, E.M. (2018). Analysis of Murine Hematopoietic Stem Cell Proliferation During Inflammation. Methods Mol Biol *1686*, 183-200. 10.1007/978-1-4939-7371-2_14.

Kanehisa, M., and Goto, S. (2000). KEGG: kyoto encyclopedia of genes and genomes. Nucleic Acids Res *28*, 27-30. 10.1093/nar/28.1.27.

Kumanogoh, A., Shikina, T., Suzuki, K., Uematsu, S., Yukawa, K., Kashiwamura, S., Tsutsui, H., Yamamoto, M., Takamatsu, H., Ko-Mitamura, E.P., et al. (2005). Nonredundant roles of Sema4A in the immune system: defective T cell priming and Th1/Th2 regulation in Sema4A-deficient mice. Immunity *22*, 305-316. 10.1016/j.immuni.2005.01.014.

Li, H., Handsaker, B., Wysoker, A., Fennell, T., Ruan, J., Homer, N., Marth, G., Abecasis, G., Durbin, R., and Genome Project Data Processing, S. (2009). The Sequence Alignment/Map format and SAMtools. Bioinformatics *25*, 2078-2079. 10.1093/bioinformatics/btp352.

Liberzon, A., Birger, C., Thorvaldsdottir, H., Ghandi, M., Mesirov, J.P., and Tamayo, P. (2015). The Molecular Signatures Database (MSigDB) hallmark gene set collection. Cell Syst *1*, 417-425. 10.1016/j.cels.2015.12.004.
Love, M.I., Huber, W., and Anders, S. (2014). Moderated estimation of fold change and dispersion for RNA-seq data with DESeq2. Genome Biol *15*, 550. 10.1186/s13059-014-0550-8.
Lun, A.T., Bach, K., and Marioni, J.C. (2016). Pooling across cells to normalize single-cell RNA sequencing data with many zero counts. Genome Biol *17*, 75. 10.1186/s13059-016-0947-7.
McInnes, L., Healy, J., and Melville, J. (2020). UMAP: Uniform Manifold Approximation and Projection for Dimension Reduction. arXiv *1802.03426*.
Merico, D., Isserlin, R., Stueker, O., Emili, A., and Bader, G.D. (2010). Enrichment map: a network-based method for gene-set enrichment visualization and interpretation. PLoS One *5*, e13984. 10.1371/journal.pone.0013984.
Okamoto, K., Uchida, S., Ito, T., and Mizuno, N. (2007). Self-organization of all-inorganic dodecatungstophosphate nanocrystallites. J Am Chem Soc *129*, 7378-7384. 10.1021/ja070694c.
Patro, R., Duggal, G., Love, M.I., Irizarry, R.A., and Kingsford, C. (2017). Salmon provides fast and bias-aware quantification of transcript expression. Nat Methods *14*, 417-419. 10.1038/nmeth.4197.
Picelli, S., Bjorklund, A.K., Faridani, O.R., Sagasser, S., Winberg, G., and Sandberg, R. (2013). Smart-seq2 for sensitive full-length transcriptome profiling in single cells. Nat Methods *10*, 1096-1098. 10.1038/nmeth.2639.
Quinlan, A.R., and Hall, I.M. (2010). BEDTools: a flexible suite of utilities for comparing genomic features. Bioinformatics *26*, 841-842. 10.1093/bioinformatics/btq033.
Scialdone, A., Natarajan, K.N., Saraiva, L.R., Proserpio, V., Teichmann, S.A., Stegle, O., Marioni, J.C., and Buettner, F. (2015). Computational assignment of cell-cycle stage from single-cell transcriptome data. Methods *85*, 54-61. 10.1016/j.ymeth.2015.06.021.
Shannon, P., Markiel, A., Ozier, O., Baliga, N.S., Wang, J.T., Ramage, D., Amin, N., Schwikowski, B., and Ideker, T. (2003). Cytoscape: a software environment for integrated models of biomolecular interaction networks. Genome Res *13*, 2498-2504. 10.1101/gr.1239303.
Sipkins, D.A., Wei, X., Wu, J.W., Runnels, J.M., Cote, D., Means, T.K., Luster, A.D., Scadden, D.T., and Lin, C.P. (2005). In vivo imaging of specialized bone marrow endothelial microdomains for tumour engraftment. Nature *435*, 969-973. 10.1038/nature03703.
Subramanian, A., Tamayo, P., Mootha, V.K., Mukherjee, S., Ebert, B.L., Gillette, M.A., Paulovich, A., Pomeroy, S.L., Golub, T.R., Lander, E.S., and Mesirov, J.P. (2005). Gene set enrichment analysis: a knowledge-based approach for interpreting genome-wide expression profiles. Proc Natl Acad Sci U S A *102*, 15545-15550. 10.1073/pnas.0506580102.
Traag, V.A., Waltman, L., and van Eck, N.J. (2019). From Louvain to Leiden: guaranteeing well-connected communities. Sci Rep *9*, 5233. 10.1038/s41598-019-41695-z.
Wolf, F.A., Angerer, P., and Theis, F.J. (2018). SCANPY: large-scale single-cell gene expression data analysis. Genome Biol *19*, 15. 10.1186/s13059-017-1382-0.
Wolf, F.A., Hamey, F.K., Plass, M., Solana, J., Dahlin, J.S., Gottgens, B., Rajewsky, N., Simon, L., and Theis, F.J. (2019). PAGA: graph abstraction reconciles clustering with trajectory inference through a topology preserving map of single cells. Genome Biol *20*, 59. 10.1186/s13059-019-1663-x.
Wolock, S.L., Lopez, R., and Klein, A.M. (2019). Scrublet: Computational Identification of Cell Doublets in Single-Cell Transcriptomic Data. Cell Syst *8*, 281-291 e289. 10.1016/j.cels.2018.11.005.
Xia, J., Swiercz, J.M., Banon-Rodriguez, I., Matkovic, I., Federico, G., Sun, T., Franz, T., Brakebusch, C.H., Kumanogoh, A., Friedel, R.H., et al. (2015). Semaphorin-Plexin Signaling Controls Mitotic Spindle Orientation during Epithelial Morphogenesis and Repair. Dev Cell *33*, 299-313. 10.1016/j.devcel.2015.02.001.

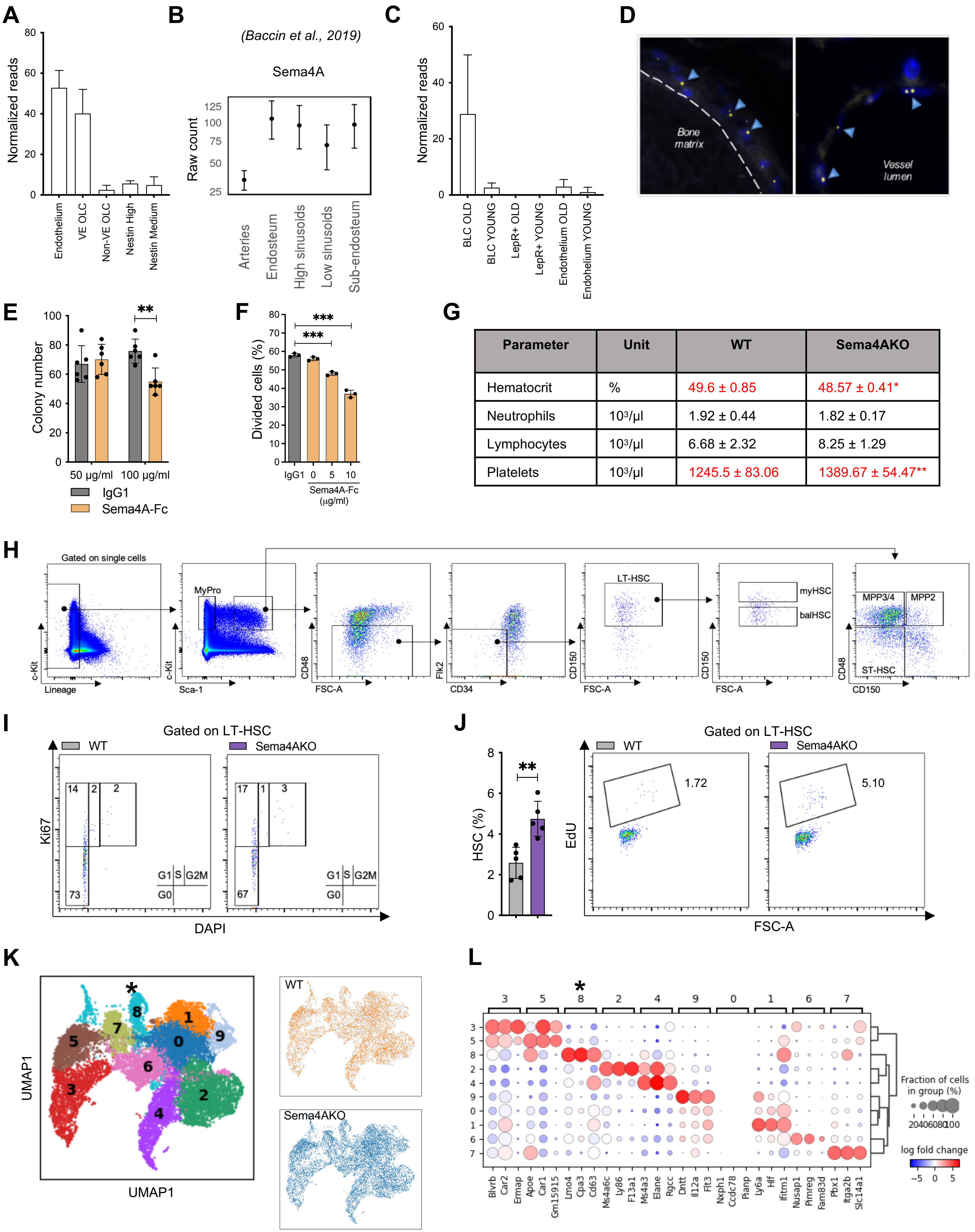


| Parameter | Unit | WT | Sema4AKO |
|---|---|---|---|
| Hematocrit | % | 49.6 ± 0.85 | 48.57 ± 0.41* |
| Neutrophils | $10^3$/µl | 1.92 ± 0.44 | 1.82 ± 0.17 |
| Lymphocytes | $10^3$/µl | 6.68 ± 2.32 | 8.25 ± 1.29 |
| Platelets | $10^3$/µl | 1245.5 ± 83.06 | 1389.67 ± 54.47** |

**Supplementary Figure 1**

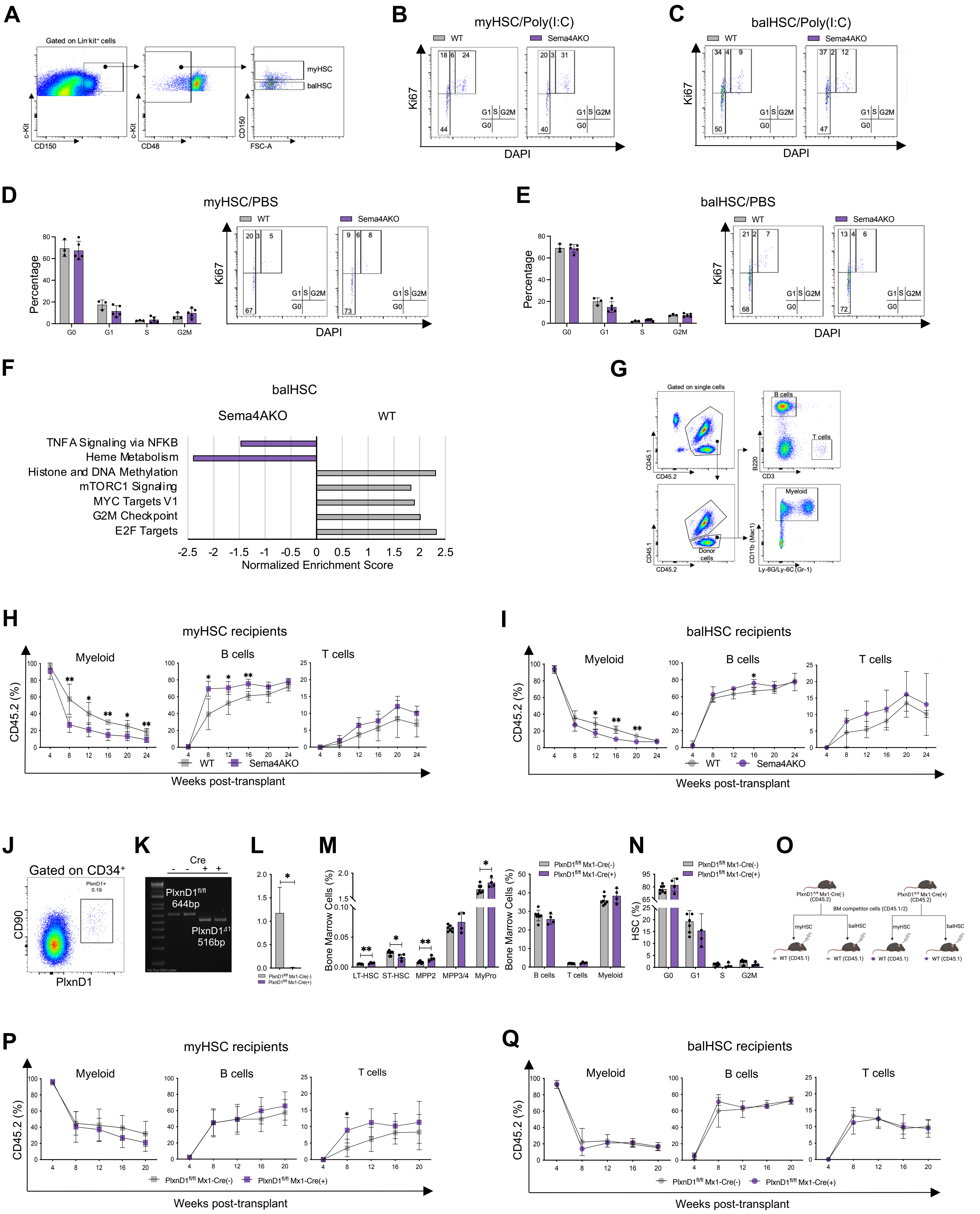


**Supplementary Figure 2**

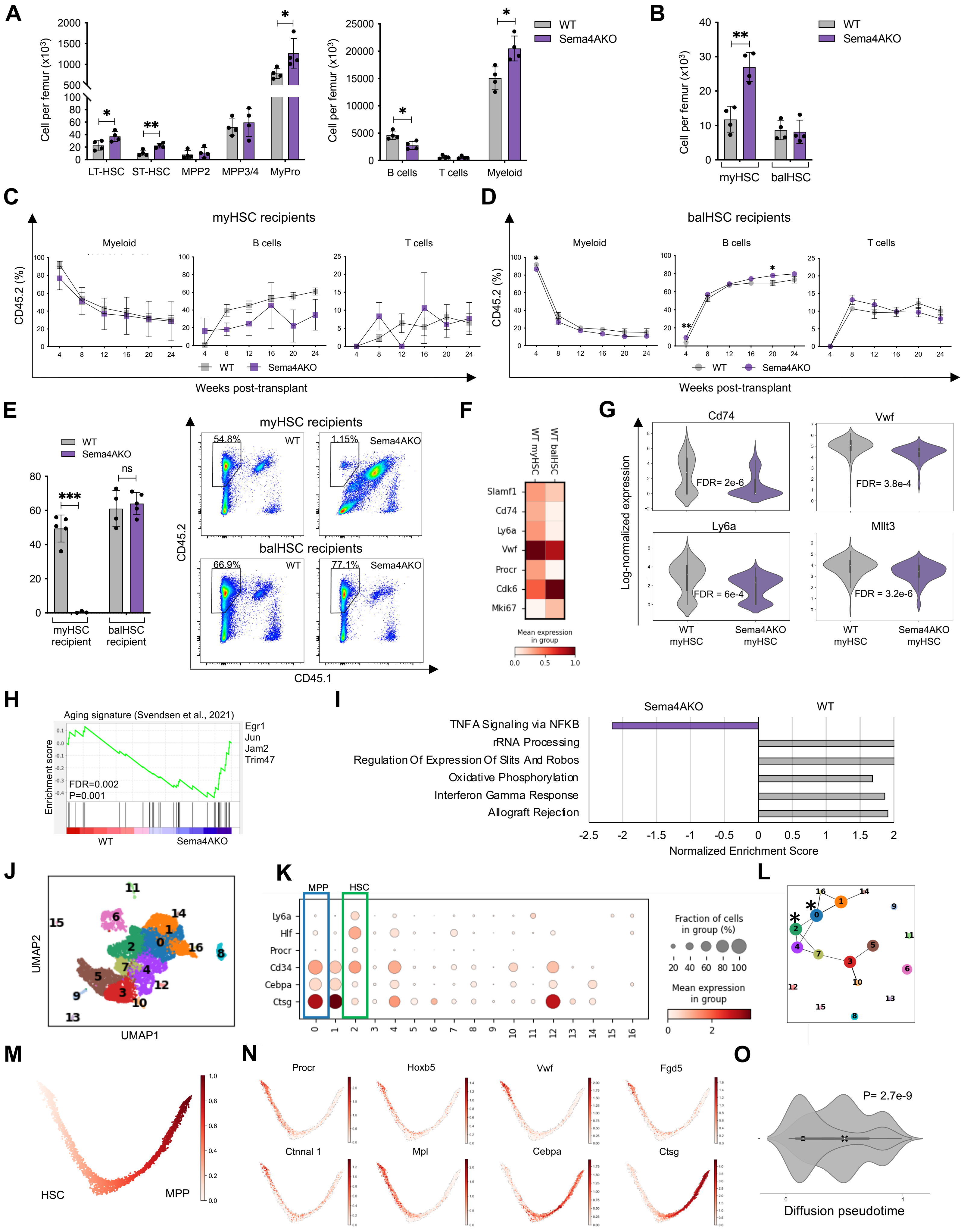


**Supplementary Figure 3**

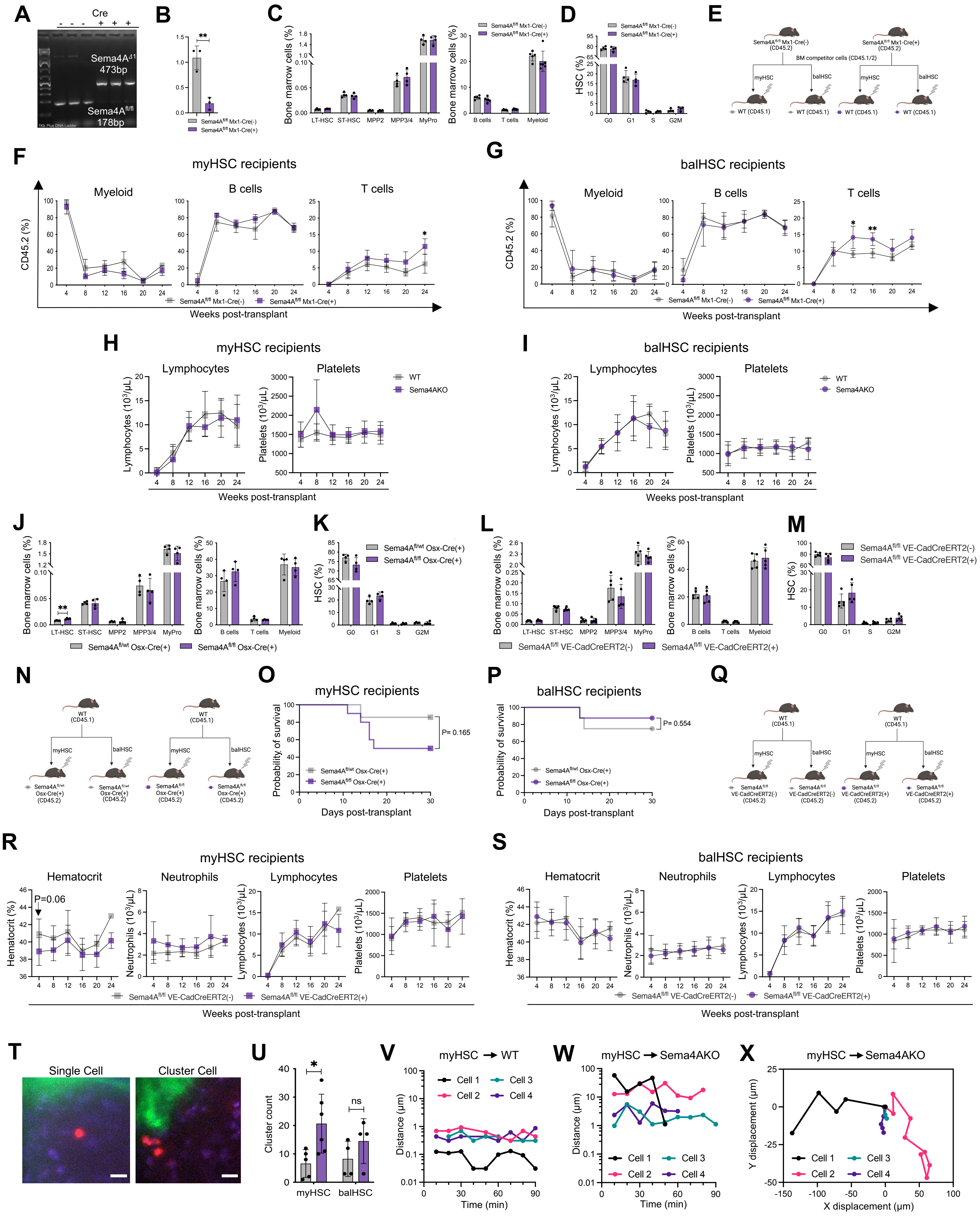


**Supplementary Figure 4**

### SUPPLEMENTARY FIGURE LEGENDS

**Figure S1. Sema4A regulates quiescence of mouse and human hematopoietic stem/progenitor cells. Related to Figure 1.**

(A) Sema4A mRNA expression in mouse niche cell subsets by bulk RNA-Seq (n=3 mice per condition).

(B) Sema4A mRNA expression in mouse niche cells by single cell RNA-Seq based on the data by Baccin et al.

(C) Sema4A mRNA expression in niche cell subsets from young vs aged WT mice (n=3 mice per condition).

(D) Human Sema4A mRNA expression in bone-lining cells (left panel) and endothelium (right panel) by single-molecule RNA FISH (RNAScope).

(E) Number of hematopoietic colonies (CFU) in the presence of mouse Sema4A-Fc/IgG1 control protein (n=6 per group).

(F) CFSE dilution analysis of *ex vivo* proliferation kinetics of human CD34+ cells 24 hours after addition of human Sema4A-Fc/IgG1 protein (n=3 technical replicates per condition, Donor 2).

(G) Complete blood count data from young WT/Sema4A KO mice (n=6 mice per genotype).

(H) Gating strategy for flow cytometric analysis/sorting of HSPC and myHSC and balHSC.

(I) Representative flow cytometry plots for cell cycle analysis of HSC from young WT/Sema4AKO mice.

(J) Short-term EdU incorporation (cumulative data and representative plots) by HSC from WT/Sema4AKO mice (n=5 mice per genotype).

(K, L): UMAP of 13417 WT (n=1) and Sema4AKO (n=2) lin-kit+ bone marrow cells with Leiden clustering. Distribution of WT (orange) and Sema4AKO (blue) cells within UMAP are shown. Cluster 1, highlighted with an asterisk, is determined to be HSC due to expression of HSC markers *Ly6a* and *Hlf* shown in L, a mean expression dot-plot of 3 markers per cluster.

Data are presented as mean ± SD *p < 0.05; **p < 0.01; ***p < 0.001 by unpaired T-test.

**Figure S2. Sema4A/PlxnD1 signaling constrains the response of myeloid-biased HSC to proliferative stress. Related to Figure 2.**

(A) Gating strategy for Poly (I:C) injection experiments.

(B) Representative myHSC cell cycle plots from Poly (I:C)-injected WT/Sema4AKO mice.

(C) Representative balHSC cell cycle plots from Poly (I:C)-injected mice WT/Sema4AKO mice.

(D) Bar graph and representative myHSC cell cycle plots from PBS-injected WT/Sema4AKO mice (n=3 mice per genotype).

(E) Bar graph and representative balHSC cell cycle plots from PBS-injected WT/Sema4AKO (n=3 mice per genotype).

(F) GSEA of balHSC from WT/Sema4AKO Poly(I:C) injected mice, FDR<0.01.

(G) Gating strategy for post-transplant chimerism analysis.

(H) Lineage composition of donor-derived cells from WT mice transplanted with WT/Sema4AKO myHSC (n=5 recipients per genotype).

(I) Lineage composition of donor-derived cells from WT mice transplanted with WT/Sema4AKO balHSC (n=4-5 recipients per genotype).

(J) Representative flow cytometry plot of PlxnD1 expression in human $CD34^{+}CD90^{+}$ cells.

(K) Representative gel photograph of PlxnD1 excision validation. PCR analysis was performed using genomic DNA from LKS cells as per indicated genotypes.

(L) PlxnD1 Q-PCR analysis of mRNA from LKS cells as per indicated genotypes.

(M) Immunophenotypic analysis of the bone marrow from $PlxnD1^{fl/fl}$ Mx1Cre(+) and $PlxnD^{fl/fl}$ Mx1Cre(-) mice (n=4-7 mice per genotype).

(N) HSC cell cycle analysis in $PlxnD1^{fl/fl}$ Mx1-Cre(+) and $PlxnD^{fl/fl}$ Mx1-Cre(-) mice (n=4-7 mice per genotype).

(O) Experimental schema for the transplant studies shown in Fig. 2K and Fig. 2L.

(P) Lineage composition of donor-derived cells in the recipients of myHSC from $PlxnD1^{fl/fl}$ Mx1-Cre(+) and $PlxnD^{fl/fl}$ Mx1-Cre(-) mice (n=5 recipient mice per donor genotype).

(Q) Lineage composition of donor-derived cells in the recipients of balHSC from $PlxnD1^{fl/fl}$ Mx1-Cre(+) and $PlxnD^{fl/fl}$ Mx1-Cre(-) mice (n=5 recipient mice per donor genotype).

Data are presented as mean ± SD *p < 0.05; **p < 0.01; ***p < 0.001 by unpaired T-test

**Figure S3. Sema4A prevents excessive myHSC expansion and functional loss with age. Related to Figure 3.**

(A) Absolute number of HSPC and mature cells in aged WT/Sema4AKO mice (n=4 mice per genotype).

(B) Absolute number of myHSC and balHSC in aged WT/Sema4AKO mice (n=4 mice per genotype).

(C) Lineage composition of donor-derived cells in the recipients of myHSC from aged WT/Sema4AKO mice (n=3-5 recipients per genotype).

(D) Lineage composition of donor-derived cells in the recipients of balHSC from aged WT/Sema4AKO mice (n=4-5 recipients per genotype).

(E) Donor-derived (CD45.2) bone marrow chimerism in the recipients of myHSC and balHSC from aged WT/Sema4AKO mice. Representative flow cytometry plots are shown on the right (n=3-5 recipients per genotype).

(F) Scaled mean expression of HSC cell cycle/self-renewal genes in WT balHSC vs WT myHSC from aged WT/Sema4AKO mice.

(G) Single cell expression (and corresponding FDR values) of HSC cell cycle/self-renewal genes which were significantly downregulated in aged Sema4AKO myHSC.

(H) GSEA plot and top differentially expressed genes for the "core aging signature" (Svendsen et al, 2021) in aged WT/Sema4AKO myHSC.

(I) GSEA of balHSC from aged WT/Sema4AKO mice, FDR<0.01.

(J) UMAP of 8531 WT (N=2) $lin^-$ $kit^+$ bone marrow cells with Leiden clustering.

(K) Dot-plots of mean expression for selected HSPC markers. HSC (cluster 2) and MPP (cluster 0) are highlighted.

(L) Direct lineage relationship between cluster 2 and cluster 4, as predicted by graph abstraction (PAGA).

(M) Diffusion map representing HSC differentiation trajectory. Cells are colored based on diffusion pseudotime (DPT) coordinates.

(N) Expression of HSC/MPP marker genes along the DPT trajectory. Cells are colored based on their normalized expression levels for each the gene indicated at the top.

(O) Distributions of diffusion pseudotime values of WT aged myHSC and balHSC. The P-value is estimated with a Wilcoxon-rank sum test.

Data are presented as mean ± SD *p < 0.05; **p < 0.01; ***p < 0.001 by unpaired T-test unless otherwise stated.

**Figure S4. Sema4A from the bone marrow niche restrains stress-induced myHSC proliferation and maintains self-renewal. Related to Figure 4.**

(A) Representative gel photograph of Sema4A excision validation. PCR analysis was performed using genomic DNA from LKS cells as per indicated genotypes.

(B) Sema4A Q-PCR analysis of mRNA from LKS cells as per indicated genotypes.

(C) Immunophenotypic analysis of the bone marrow from $Sema4A^{fl/fl}$ Mx-1Cre(+) and $Sema4A^{fl/fl}$ Mx1-Cre(-) mice (n=4-6 mice per genotype).

(D) HSC cell cycle analysis in $Sema4A^{fl/fl}$ Mx-1Cre(+) and $Sema4A^{fl/fl}$ Mx1-Cre(-) mice (n=4 mice per genotype).

(E) Experimental schema for the transplantation experiment shown in F and G.

(F) Lineage composition of donor-derived cells in the recipients of myHSC from Sema4A$^{fl/fl}$ Mx1-Cre(+) and Sema4A$^{fl/fl}$ Mx1-Cre(-) mice (n=4-5 recipients per genotype).

(G) Lineage composition of donor-derived cells in the recipients of from Sema4A$^{fl/fl}$ Mx1-Cre(+) and Sema4A$^{fl/fl}$ Mx1-Cre(-) mice (n=4-5 recipients per genotype).

(H, I) Post-transplant lymphocyte and platelet count for WT/Sema4AKO recipients of myHSC (panel H) and balHSC (panel I) (n=9-11 recipients per genotype).

(J) Immunophenotypic analysis of the bone marrow from Sema4A$^{fl/fl}$ Osx-Cre(+) and Sema4A$^{fl/wt}$ Osx-Cre(+) mice (n=4).

(K) HSC cell cycle analysis in Sema4A$^{fl/fl}$ Osx-Cre(+) and Sema4A$^{fl/wt}$ Osx-Cre(+) mice (n=4 mice per genotype).

(L) Immunophenotypic analysis of the bone marrow from Sema4A$^{fl/fl}$ VE-CadCre ERT2(+) and Sema4A$^{fl/fl}$ VE-CadCre ERT2(-) mice (n=5).

(M) HSC cell cycle analysis in Sema4A$^{fl/fl}$ VE-CadCre ERT2(+) and Sema4A$^{fl/fl}$ VE-CadCreERT2(-) mice (n=5 mice per genotype).

(N) Experimental schema for the transplantation experiment using Sema4A$^{fl/fl}$ Osx-Cre (+) and Sema4A$^{fl/wt}$ Osx-Cre(+) mice.

(O, P) Survival curves from the experiments depicted in (N) (n=7-10 recipients per genotype in each group).

(Q) Experimental schema for the transplantation experiment using Sema4A$^{fl/fl}$ VE-CadCre ERT2(+) and Sema4A$^{fl/fl}$ VE-CadCre ERT2(-) recipient mice.

(R,S) Post-transplant blood counts from experiments depicted in R (n=5-7 recipients per genotype in each group).

(T) Representative two-photon intravital images of DiD-labeled single cells and clusters in the calvarial bone marrow of transplanted mice. Cells (DiD, red), bone (SHG, green), and autofluorescence (blue) are shown. Scale bars ~25 μm.

(U) Average number of clusters per mouse ~15-20 hours after transplantation of WT myHSC or balHSC into WT/Sema4AKO recipients, as assessed by two-photon intravital imaging of the calvarial bone marrow (n = 33, 124, 33 and 58 cell clusters for WT myHSC, Sema4AKO myHSC, WT balHSC, and Sema4AKO balHSC, respectively; data are the summary of 6 independent experiments involving a total of n = 4-6 mice per group).

(V-W) Cell displacement over time determined by two-photon time-lapse imaging of WT myHSC transplanted into WT (panel V) and Sema4AKO (panel W) recipients, respectively (data are the summary of 6 independent experiments involving a total of n = 5-6 mice per group). These 4 myHSC in Sema4AKO recipients were the only cells found to be motile over all imaging experiments for all mice. Some motile cells either entered the FOV, exited the FOV or became immobile part way through imaging. The 4 myHSC in WT recipients are representative non-motile cells and had a total displacement < 2 μm over 1.5 hours of imaging.

(X) X/Y cell displacement of motile WT myHSC transplanted into Sema4AKO mice determined by two-photon time-lapse imaging of the calvaria (data are the summary of 6 independent experiments involving a total of n = 6 mice).

Data are presented as mean ± SD *$p < 0.05$; **$p < 0.01$; ***$p < 0.001$ by unpaired T-test.
